# The developmental progression of eight opsin spectral signals recorded from the zebrafish retinal cone layer is altered by the timing and cell type expression of thyroxin receptor β2 (trβ2) gain-of-function transgenes

**DOI:** 10.1101/2022.06.02.494548

**Authors:** Ralph F. Nelson, Annika Balraj, Tara Suresh, Leah J. Elias, Takeshi Yoshimatsu, Sara S. Patterson

**Affiliations:** National Institute of Neurological Disorders and Stroke, National Institutes of Health, Rockville, Maryland 20892; Department of Biological Structure, University of Washington, Seattle, Washington 98195

## Abstract

Zebrafish retinal cone signals shift in spectral shape through larval, juvenile, and adult development as expression patterns of eight cone-opsin genes change. An algorithm extracting signal amplitudes for the component cone spectral types is developed and tested on two thyroxin receptor β2 (trβ2) gain-of-function lines *crx:mYFP-2A-trβ2* and *gnat2:mYFP-2A-trβ2*, allowing correlation between opsin signaling and opsin immunoreactivity in lines with different developmental timing and cell-type expression of this red-opsin-promoting transgene. Both adult transgenics became complete, or nearly complete, ‘red-cone dichromats’, with disproportionately large LWS1 opsin amplitudes as compared to controls, where LWS1 and LWS2 amplitudes were about equal, and significant signals from SWS1, SWS2, and Rh2 opsins were detected. But in transgenic larvae and juveniles of both lines it was LWS2 amplitudes that increased, with LWS1 cone signals rarely encountered. In *gnat2:mYFP-2A-trβ2* embryos at 5 days post fertilization (dpf), red-opsin immunoreactive cone density doubled, but red-opsin amplitudes (LWS2) increased < 10%, and green, blue and UV opsin signals were unchanged, despite co-expressed red opsins, and the finding that an *sws1* UV-opsin reporter gene was shut down by the *gnat2:mYFP-2A-trβ2* transgene. By contrast both LWS2 red-cone amplitudes and the density of red-cone immunoreactivity more than doubled in 5 dpf *crx:mYFP-2A-trß2* embryos, while UV-cone amplitudes were reduced 90%. Embryonic cones with trβ2 gain-of-function transgenes were morphologically distinct from control red, blue or UV cones, with wider inner segments and shorter axons than red cones, suggesting cone spectral specification, opsin immunoreactivity and shape are influenced by the abundance and developmental timing of trβ2 expression.

**Significance Statement:** As different combinations of eight cone-opsin mRNAs are successively expressed during zebrafish development and maturation, the composite cone-ERG spectral signal shifts. Amplitudes of each of the eight resulting cone signals are inferred computationally from the composite signal, both in controls and in two thyroxin-receptor β2 (trβ2) gain-of-function transgenics, *crx:mYFP-2A-trβ2* and *gnat2:mYFP-2A-trβ2*, trβ2 being a transcription factor required for expression of the red-cone opsins LWS1 and LWS2. Adult transgenics become red cone dichromats with excess LWS1 amplitudes, but larvae and juveniles evoke excess LWS2 amplitudes. Controls retain 5 to 6 cone signals of changing composition throughout development. The progression of transgene-induced amplitude alterations is slower in *gnat2:mYFP-2A-trβ2*, with supernormal red-opsin antigenicity not immediately correlating with red-cone signaling.

## Introduction

Spectral patterns in the zebrafish cone ERG shift with development (Saszik et al., 1999; Nelson et al., 2019). These shifts are determined in part by factors regulating opsin expression. We develop an algorithm to extract the electrical contributions of each of the eight zebrafish cone opsins from massed ERG cone signals and use this tool to examine alterations in cone-signal development brought about by perturbations in expression of the regulatory factor thyroid hormone receptor β2 (trß2). The process examines the correlation of opsin expression with opsin signal development. Thyroid hormone receptor β2, a splice variant of the *thrb* gene, is selectively expressed in vertebrate retinal cones (Ng et al., 2001). When deleted, cones expressing opsins in the long-wavelength-sensitive (LWS) subfamily of the opsin molecular phylogenetic tree (Terakita, 2005) fail to be produced. This includes the MWS cones of mouse (Ng et al., 2001; Roberts et al., 2006), human MWS and LWS cone function (Weiss et al., 2012), and both LWS1 and LWS2 red-cone signals of zebrafish (Deveau et al., 2020). The LWS cone subfamily senses the longest wavelengths a species detects, with opsin spectral peaks ranging from 511 nm in mouse (Jacobs et al., 1991) to 625 nm in goldfish (Marks, 1965).

Two gain-of-function transgenics, *crx:mYFP-2A-trβ2* (*crx:trβ2*) and *gnat2:mYFP-2A-trß2;mpvl7^-/-^* (*gnat2:trβ2*) (Suzuki et al., 2013), are used to perturb both the developmental timing and the cellular locus of trß2 expression. The first (*crx:trβ2*) is active by day 2 in larval development, in embryonic retinal progenitor cells (Shen and Raymond, 2004; Suzuki et al., 2013). In the second (*gnat2:trβ2*), trβ2 is promoted by *gnat2*, the cone transducin α promoter (Brockerhoff et al., 2003; Kennedy et al., 2007). This introduces trβ2 later in development, and only into differentiated retinal cones. In each transgenic Suzuki et al (2013) showed an excess of red-opsin immunoreactive cones. In *crx:trβ2* the larval densities of green-, blue-, and UV -opsin immunoreactive cones were reduced, whereas in *gnat2:trβ2* excess red-opsin immunoreactivity found a home in cones expressing other opsins, forming mixed-opsin cones. These transgenic mixed-opsin cones were thought to model rodent cones that natively express both UV and MWS opsins (Applebury et al., 2000). We here confirm the immunoreactive patterns of these transgenics under the same conditions used for electrophysiological recordings of the signals from individual cone types to determine whether altered opsin patterns have a parallel in cone spectral signals. The physiological consequences of altered trβ2 expression are unknown, and might result in either retinal disease, or a spectrally unique visual system.

Although trβ2 might be thought of as a binary ON/OFF switch for LWS-cone development, it clearly has other actions, and the activity level and cell-type expression may be important. Dilution of trβ2 in adult zebrafish *trβ2*^+/-^ animals made red cones less dense and depressed long-wavelength sensitivity (Deveau et al., 2020). In embryonic and juvenile zebrafish, while unbound trβ2 sufficed for red-cone development, binding of exogenous thyroid hormone (TH) shifted larval cones expressing LWS2 opsin to expression of LWS1 opsin, and athyroidism switched them back (Mackin et al., 2019). TH depressed the expression of both SWS1- (UV-opsin) and SWS2- (blue-opsin) message (Mackin et al., 2019). TH and trβ2 appear further to be involved in the preferential expression of several of the tandem quadruplicate Rh2 cone opsins in zebrafish (Mackin et al., 2019). It appears likely that TH and trβ2 are upstream regulators in the specification of all cone types, though only for red cone neuronal specification is trβ2 essential (Deveau et al., 2020). Therefore, the developmental impact on electrical signaling in all eight zebrafish cone types is investigated.

## Materials and Methods

### Zebrafish (Danio rerio) husbandry and experimental design

The transgenic gain-of-function and reporter-line zebrafish *crx:mYFP-2A-trβ2, gnat2:mYFP-2A-trβ2;mpvl7^-/-^, trβ2:tdTomato;sws1:GFP, sws1:nfsBmCherry* and *sws2:GFP* (Takechi et al., 2003; Takechi et al., 2008; Suzuki et al., 2013) were generously provided by the Rachel Wong lab (University of Washington). In gain-of-function studies, larvae were obtained by outcrosses such that transgenics were heterozygotes and controls were siblings (Fig. 1A), studied at random on the same developmental days post fertilization. The transgenic *gnat:mYFP-2A-trβ2* was maintained on an *mpv17^-/-^* (*roy orbison*) background (D’Agati et al., 2017) (Fig. 1C). There are several studied mutations of *mpv17* all with loss of irridophores (Bian et al., 2021). Here we use the naturally occurring 19-base-pair intronic deletion of the *roy orbison* mutant (D’Agati et al., 2017). The lack of reflective iridophores in the iris of this mutant facilitated phenotyping of pupil fluorescence. Pineal fluorescence was also a *gnat:mYFP-2A-trβ2;mpv17*characteristic.

**Figure 1.**
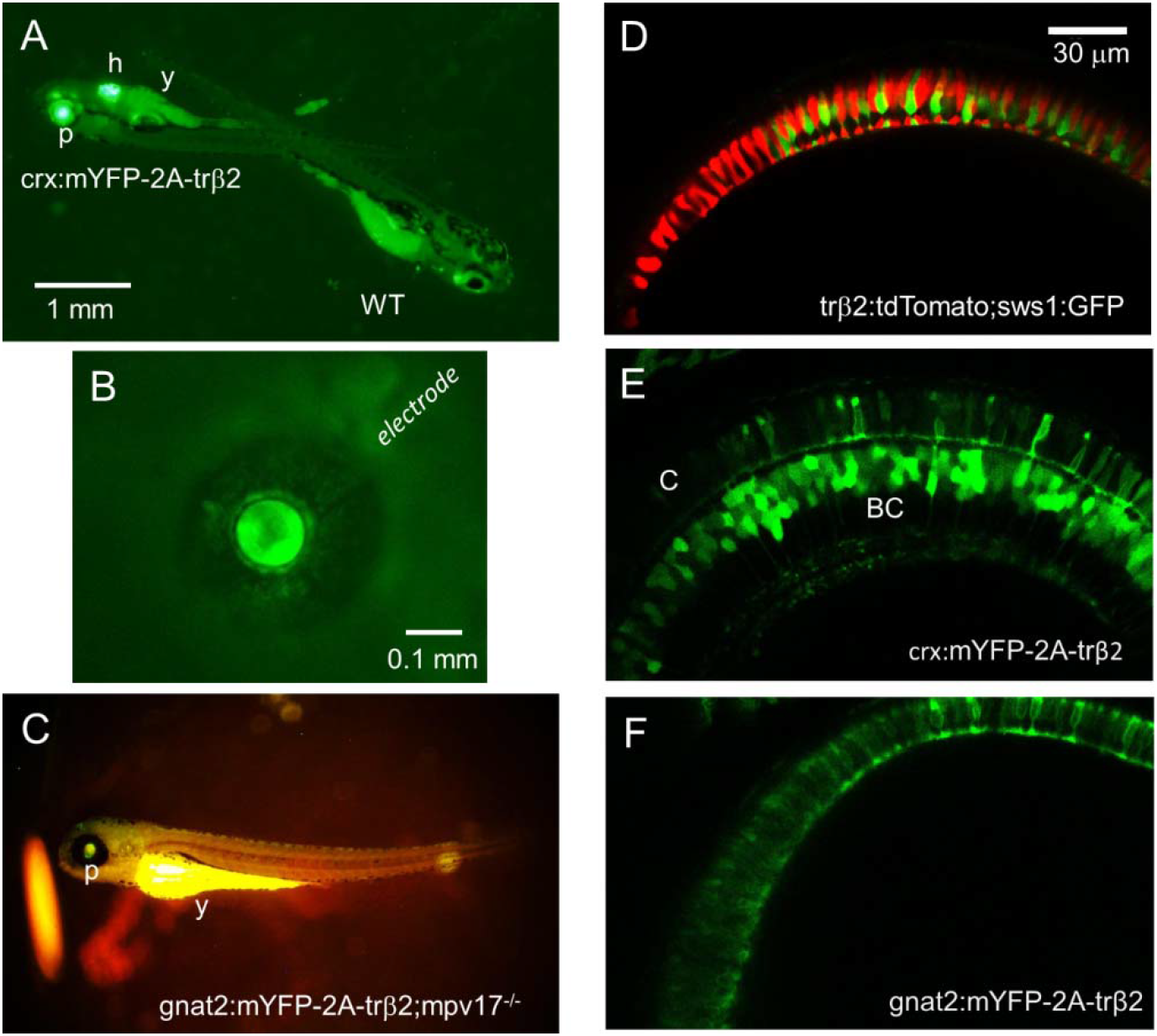
Reporter fluorescence of zebrafish larvae. ***A***, Larvae from outcrosses of *crx:mYFP-2A-trβ2* heterozygotes are either heterozygous or wild-type controls (WT). The heterozygotes are recognized by both pupil (p) and heart (h) fluorescence. The yolk sac (y) is autofluorescent. ***B***, The cornea of a 5-day transgenic eye isolated from a *crx:mYFP-2A-trβ2* larva is penetrated with a patch electrode for ERG recordings. The eye is less than 0.5 mm diameter. ***C***, The *gnat2:mYFP-2A-trβ2;mpv17^-/-^* gain-of-function phenotype is studied on a *roy orbison (mpv17^-/-^*) background strain. The darkly pigmented, non-reflective iris of this control strain aids in visualizing the dim transgenic fluorescence of the pupil (p). The yoke (y) is autofluorescent. (***D***, ***E***, ***F***) Live confocal imaging of retinas in 6 dpf larvae.***D***, WT red (red) and UV (green) cone morphology visualized with *trβ2:tdTomato* and *sws1:GFP* fluorescent reporter transgenes. ***E***, The *mYFP* construct in the *crx:mYFP-2A-trβ2* transgene causes cones (C) and bipolar cells (BC) to fluoresce. ***F***, The *mYFP* construct in the transgene *gnat2:mYFP-2A-trβ2* marks only cone cells.

Spectral physiology was collected from larvae on days 5, 6, 7 days post fertilization (dpf), on juveniles (12 dpf), and a from both male and female adults (8-18 mo). Larvae lack gender. For 5-7 dpf, larvae were group incubated at 28°C in 3.5-inch Petri dishes atop a heating pad. Larval medium contained 60 mg/liter sea salt, 75 μl/liter 1.5% methylene blue (Sigma-Aldrich Cat. No. 03978). 5-7 dpf larvae were not fed. The methylene blue was omitted for live confocal microscopy because of its fluorescence. 8-12-day larvae were group housed in system nursery tanks (520-650 μΩwater, 28°C, pH 7.5-7.7) on the same light/dark cycle, and fed both Larval AP100 (Pentair Aquatic Eco-Systems) and live rotifers (*Brachionus plicatilis;* Reed Mariculture).

Some larvae were raised to adulthood (8-18 months) both for spectral studies and for retention as breeders. These were group housed using 1.5- or 3-liter recirculating tanks shelved in stand-alone, recirculating, Aquatic Habitats benchtop systems (Pentair Aquatic Eco-Systems, Apopka, Florida) on the facility 14/10-hour light/dark cycle (ZT0 = 8:00 AM). Adults were fed pulverized TetraMin flakes (Tetra GMBH) and live rotifers. The protocols for breeding and experimentation were approved by the National Institute of Neurological Disorders and Stroke / National Institute on Deafness and Other Communication Disorders / National Center for Complementary and Integrative Health IACUC (ASPs 1307, 1227).

### Preparation and perfusion of Isolated eyes for physiology

Larvae were captured with disposable pipettes and placed on a glass lantern slide. After removing excess water, larvae were adsorbed onto a chip of black nitrocellulose filter paper (Millipore, 0.45 mm pore, Cat. No. HABP02500, MilliporeSigma), and decapitated (without anesthetic) using a long (37 mm) insect pin (Carolina Biological Supply). Using a binocular microscope (MZ12-5; Leica Microsystems), a longitudinal, dorsal-ventral cut through the head proptosed and isolated larval eyes, which were positioned facing up, taking care not to touch the eye directly. In the recording chamber, larval eyes mounted on the nitrocellulose chip were perfused at 0.1 ml/min with Minimal Essential Medium (MEM; Thermo Fisher Scientific Cat. No. 11090-099, equilibrated with 95% O_2_ and 5% CO_2_) using a syringe pump (New Era 500L; Braintree Scientific) and a 28-guage microfil syringe needle as applicator (MF28G67; World Precision Instruments). The chamber was an inverted lid for a 35-mm culture dish (ThermoFisher Scientific), with a disk of 41-μm-mesh nylon filter (Millipore) covering the bottom to wick away perfusate. The perfusion applicator was positioned on the nylon mesh. 20 mM L-Aspartate (Sigma-Aldrich), added to the MEM perfusate, blocked post-synaptic, glutamatergic, photoreceptor mechanisms leaving only photoreceptor signals (Sillman et al., 1969). Aspartate medium blocks cone synaptic transmission through saturation of glutamatergic receptors of three types: ionotropic cation channels, metabotropic-mediated cation channels, and glutamate-transporter-mediated anionic channels (Grant and Dowling, 1995, 1996; Connaughton and Nelson, 2000). Patch electrodes (3 μm tip) were inserted trans-corneally (Fig. 1b) to record the isolated cone ERG signals (cone PIII) (Wong et al., 2004).

Adult eyecups were prepared from eyes removed from fish decapitated with a fresh, single-edged razor. Corneas and lenses were removed from the isolated eyes mounted upright on a 5-10 mm square of black nitrocellulose paper, and the preparation was placed in a recording chamber (as above). The perfusion applicator was placed directly above the retina oxygenating the vitreal surface with MEM containing 10 mM L-Aspartate at 0.3 ml/min. Microelectrodes broken to 300 μm tip diameter placed in the eyecup recorded cone-PIII ERG signals (Nelson and Singla, 2009).

### Live imaging of larval retinas

Transgenic larvae were raised at 28°C in 300 μM Phenylthiourea (PTU, Sigma-Aldrich) to prevent melanin formation in the pigment epithelium and allow imaging of the retina *in vivo*. At 6 dpf, each larva was mounted individually in 1.5% agarose (Sigma-Aldrich type VII-A) on an 8-chamber slide with the right eye against the cover-glass floor of the chamber. Eyes were imaged in z-stacks on Zeiss 880 confocal microscope at either 25x or 40x magnification at 1024-×-1024-pixel resolution. Cone morphometrics were measured for 5 or 6 fish in each transgenic line in Fiji (ImageJ) on the optical slice that offered the longest stretch of resolved cones. These measurements were analyzed for differences using a one-way ANOVA and Tukey’s post-hoc test. Fluorescent reporters identified the morphology of wild type (WT) red and UV cones in eyes from *trβ2:tdTomato;sws1:GFP* (Fig. 1D) and of transgenic *gnat:mYFP-2A-trβ2* cones (Fig. 1F). In *crx:mYFP-2A-trβ2*, reporter fluorescence appeared in both cones and bipolar cells (Fig. 1E) as previously seen in antibody staining for Crx in zebrafish (Shen and Raymond, 2004).

### Immunohistochemistry

Larvae were euthanized by icing, and then fixed in 4% paraformaldehyde (PFA) in 0.1M phosphate-buffered saline (PBS), pH 7.4, for 25 min at room temperature. Retinas were dissected in PBS using a pair of 30-gauge syringe needles, blocked in a solution containing 5% normal donkey serum and phosphate-buffered Triton X-100 0.5% (PBT) for 1-24 h and then incubated with primary antibodies. Triton X-100 was added to enhance antibody penetration. Primary antibodies included anti-ultraviolet opsin (rabbit, 1:500, kindly provided by David Hyde), anti-ultraviolet opsin (rabbit, 1:5,000, kindly provided by Jeremy Nathans), anti-blue opsin (rabbit, 1:5,000, kindly provided by Jeremy Nathans) and anti-rod opsin (1D4, mouse, 1:100, Santa Cruz: sc-57432, RRID:AB_785511), the later raised against bovine rhodopsin but recognizing red-cone outer segments in zebrafish (Yin et al., 2012). These were diluted into the blocking solution. After incubating for 4 days at 4 °C, samples were washed three times, 15 min each, in PBT and incubated for 1 day with secondary antibodies (DyLight 649 donkey anti-mouse, and Alexa 594 donkey anti-rabbit, RRID:AB_2340621, 1:500 each, Jackson ImmunoResearch) diluted in blocking solution. After three, 15 min washes (PBT), immunostained retinas were mounted in 0.5% agarose and cover slipped with Vectashield mounting medium (Vector, RRID_2336789). Confocal image stacks were acquired on a Leica SP8 microscope using a 1.0 NA 63 X oil-immersion objective lens. Images were typically acquired with an XY resolution of 0.077 μm per pixel and 0.25 μm-thick Z slices (Yoshimatsu et al., 2014).

### Spectral stimulus protocol

Larval eyes and adult eyecups were stimulated with nine wavelengths ranging from 330 nm to 650 nm (20-nm half-width interference filters, 40-nm increments, Chroma Technology). Seven intensities were presented at each wavelength (UV compliant neutral density filters, 0.5 log unit increments covering 3 log units, Andover Corporation). A calibrated photodiode (Newport Corporation) was used to determine stimulus irradiance in quantal units. This was placed in the plane of the cornea for larval eyes or the plane of the retina for adult eyecups. All spectral model calculations are based on absolute, wavelength-specific photodiode calibrations of quanta delivered to the eye. The light source was a 150W OFR Xenon arc with two optical channels gated by Uniblitz shutters (Vincent Associates). The stimulus channel passed through three Sutter filter wheels, through a UV-visible compliant liquid light guide (Sutter Instruments), through the epifluorescence port of the BX51WI upright microscope (Olympus –Life Science Solutions), and through either a 10 × UPlanFLN/0.3 microscope objective (larvae) or a 4 × UPlanSApo/0.16 objective (adults). The second optical channel passed through hand-inserted filters and an infrared compliant liquid light guide (Newport Corporation) providing infrared (IR) side illumination for visualization and ‘neutral’ backgrounds, or through a 627 nm interference filter for red adapting backgrounds.

The spectral protocol was a fixed sequence of 280, 300 msec, monochromatic light flashes of different irradiances and wavelengths delivered by computer using in house software. Among these stimuli were 64 unique irradiance-wavelength combinations and 6 replicates delivered 10 minutes apart as an amplitude-stability check during the 20-min protocol. The 280 stimuli created a set of 70, 4X-averaged, electroretinogram (ERG) responses, the ‘spectral dataset’. The interval between stimuli varied between 2.5 and 6 s, with the longer intervals separating the brighter stimuli. The maximal irradiances in log(quanta·μm^-2^·s^-1^) in the stimulus protocol were 7.2 (650 nm), 6.3 (610 nm), 6.4 (570 nm), 6.3 (530 nm), 6.4 (490 nm), 6.1 (450 nm), 5.7 (410 nm), 5.7 (370 nm), and 5.2 (330 nm).

To record a spectral dataset, the stimulating objective was positioned over the eye with a translation stage (MT-800; Sutter Instrument). Microelectrodes were inserted into eyes or eyecups with a micro-positioner (Sutter Instrument, MPC-385). ERG signals from the microelectrode were amplified by 10,000 (World Precision Instruments, DAM80, 0.1 Hz-1k Hz bandpass), and digitized (2000 Hz) with an Axon instruments 1440A (Molecular Devices) using Clampex 10 software. Setting the Clampex averaging feature to retain all the elements of an average, the 280 ERGs within a single spectral dataset were captured in a single file.

### Analysis of ERG signals

Datasets were imported into Origin analysis software and processed using Origin LabTalk scripts (Origin, various versions; Originlab Corporation). The 4 replicate-waveforms at each of the 70 wavelength and irradiance combinations were averaged and boxcar filtered (17 ms, one 60 Hz line-frequency cycle). Peak-to-trough amplitudes were extracted during the 850 ms following stimulus onset, an interval including both the hyperpolarizing trough and repolarizing peak of the Aspartate-isolated cone-PIII response. There was no b-wave component in this signal (Nelson and Singla, 2009). Each amplitude was associated in Origin with the wavelengths and irradiances of the stimuli, providing 70 wavelength, irradiance, and amplitude data points for each spectral dataset. Datasets with unstable response amplitudes over the collection period were rejected. For each genetic variant, multiple datasets from ~10 eyes were normalized to the maximal response of each dataset to form a cumulative dataset, which was fit to a spectral algorithm. The normalization weighted the individual datasets making up the cumulative dataset equally. Combining datasets from many eyes separated the trends in genetic alterations of spectral properties from the variations among individual eyes. Non-linear fits of models to spectral datasets used the Levenberg-Marquardt iteration algorithm provided by Origin. The algorithm finds the best spectral model and extracts amplitudes of various cone signals together with standard errors of estimate (SEs) from the cumulative cone-PIII response.

### Statistical analysis

We use *t*-tests, *F*-tests and ANOVAs to compare results from different treatments or measurements. To determine the most appropriate spectral models, *F*-tests on residual variances for different spectral models were employed. For statistical tests Graphpad Prism software (RRID:SCR_002798), web-based calculators, and statistical functions addressable within Originlabs Labtalk software were used.

### Finding the cone combinations best representing spectral datasets

Eight cone opsins are expressed in zebrafish retina (Chinen et al., 2003). The algorithm to identify the opsin signals generating the ERG spectral shape is based on the axiom that the tiny radial photocurrents of individual cones sum linearly in extracellular space to produce a net extracellular current which, by traveling through extracellular resistivity, generates an ERG photovoltage. We assume individual cone photocurrents relate to irradiance through Hill functions of exponent 1.0 and semi-saturation irradiance σ(Baylor and Fuortes, 1970) and that σ varies with wavelength in inverse proportion to each opsin absorbance. This scheme is represented in figure 2A (Equation 1). *V* is the net summed photovoltage in the cone ERG (cone PIII) which depends on *I* the stimulus irradiance in quanta and *wl* the wavelength. The *Vm_i_* values are non-linear fit values, the maximal or saturation voltages for each of the *i* cone types. The semi-saturation irradiance for the *i_th_* cone is *k_i_* as evaluated at the *i_th_* cone absorbance maximum. The *k_i_* values differ among cone types relative to each other. The relative *k_i_* values, expressed as log(*k_i_*) relative to UV-cone sensitivity, are literature values (Nelson et al., 2019) listed in figure 2C. The UV-cone value for log(*k_i_*) is fit by the algorithm. *A(wlmax_i_, wl*) is absorbance as a function of wavelength (*wl*) for the *i_th_* cone, whose wavelength maximum is *wlmax_i_*. The maximum wavelengths are literature values (Nelson et al., 2019). The Dartnall nomogram (Dartnall, 1953) is used as the absorbance function *A(wlmax_i_ wl*). This approximation has the convenience of making opsin absorbance across wavelengths a function of a single variable, *wlmax_i_*. The Dartnall nomogram posits that opsin absorbance shape is constant when plotted on a reciprocal wavelength axis. The template nomogram functions derive from suction electrode recordings of cones in *Danio aequipinnatus* (Palacios et al., 1996), a relative of zebrafish (*Danio rerio*). These templates are represented as 8^th^ order polynomials. We use a single template polynomial (Hughes et al., 1998) for all red, green, and blue cones listed in figure 2C, but a separate polynomial for UV cones (Palacios et al., 1996). The resulting absorbance shapes appear in figure 2B. Altogether there are nine parameters fit by the algorithm: eight *Vm_i_* values, and a single *k_i_* value (*k_UV_*).

**Figure 2.**
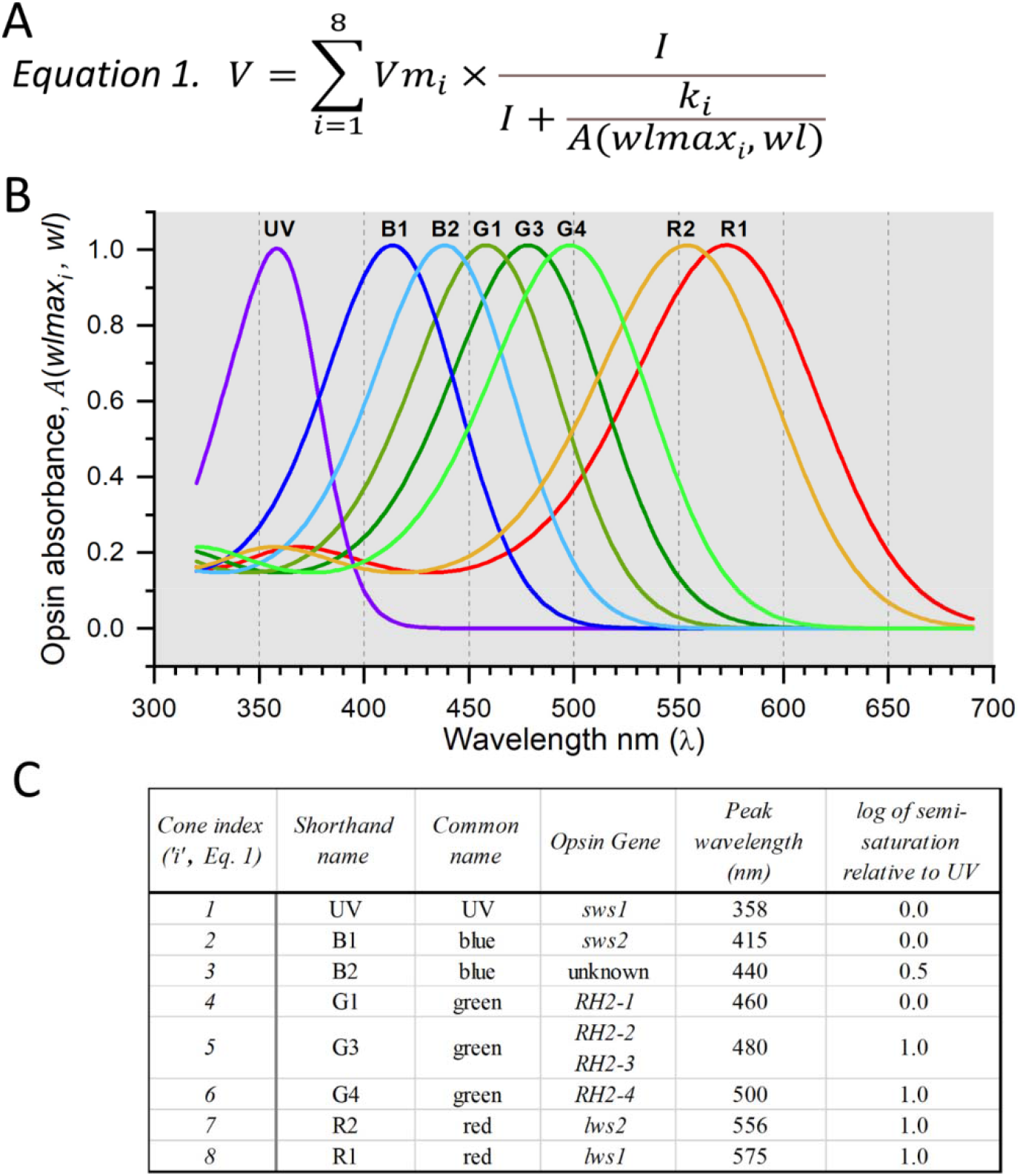
Algorithm for determining the signal strengths of the different cone types contributing to ERG spectral data. ***A***, Equation 1. Aspartate-isolated cone signals from zebrafish eyes are a summation of signals from 8 cone types, each distinguished by different maximal amplitudes (*Vm_i_*) semi-saturation irradiances (*k_i_*), and opsin peak absorbances (*wlmax_i_*). There are 255 unique combinations of 8 cones, each combination is a candidate to best model cone spectral responses in the ERG. ***B***, Spectral shapes of opsin absorbances [*A(wlmax_i_, wl*] are generated from 8^th^-order template polynomials (Palacios et al., 1996; Hughes et al., 1998) using Dartnall nomogram translations along the wavelength axis to represent opsins of different peak wavelengths (Dartnall, 1953). ***C***, Parameters for each of the *i* cone types are numbered in order from short to long wavelengths. SWS, short-wavelength-sensitive opsins; RH2, rhodopsin-like green-cone opsins; LWS, long-wavelength-sensitive opsins.

The spectral algorithm (Fig. 2A) sums signals from 8 cone types (Fig. 2B, 2C). These include a UV cone, two blue cones (B1, B2), three green cones (G1, G3, G4) and two red cones (R1, R2). The gene equivalencies of this nomenclature and the functionally measured wavelength peaks appear in figure 2C. The spectral algorithm chooses among 2^8^-1 or 255 unique cone combinations that might best represent a cumulative ERG spectral dataset. Each cone combination is called a model. The best model is chosen based on four constraints: 1) The model iteration must converge. 2) All model *Vm_i_* values must be significantly greater than zero (*t*-test, *p* ≤ 0.05). 3) All *Vm_i_* values must be less than 2.0, so as not to greatly exceed the largest amplitudes in the cumulative datasets. 4) The value of *r^2^* for the fit must be larger than that of any other model, as determined by the F-test for residual variance. If equivalent models are found (*F*-test,*p* ≥ 0.95), they will be noted. A similar modeling algorithm has been used to determine the cone combinations impinging on larval ganglion cell impulse discharges (Connaughton and Nelson, 2021).

## Results

### Cone distributions in the larval retinas of wild-type and crx:trβ2 transgenics

Suzuki et al (2013) developed the *crx:mYFP-2A-trβ2* gain-of function transgenic as a rescue line to restore red-cones during a morpholino blockade of native *trβ2* that abolished their formation. In zebrafish the *crx* gene promotor becomes activate at the retinal progenitor stage (~2 dpf) (Shen and Raymond, 2004). The gain-of-function *crx:trβ2* transgene replaces the missing retinal trβ2, but not in the same cell types or at the same developmental stage. In the experiments of Suzuki et al (2013) it was nonetheless effective. Red-opsin immunoreactive cones were restored with supernormal density, but there was suppression of green-, blue- and UV-opsin immunoreactive cones (Suzuki et al., 2013).

In figure 3A and 3B the experiment of Suzuki et al (2013) is repeated on *crx:mYFP-2A-trβ2* larvae without morpholino suppression of native *trβ2*. In WT retinas, immunoreactive mosaics of both UV cones (Fig. 3Ai) and red cones (Fig. 3Aiii) stain with opsin antibodies. Superposition of both mosaics (Fig.3Aii) show no opsin overlaps, with each cone type expressing a single opsin. The situation is similar for blue cones, which exist in a mosaic pattern separate from red cones (Figs. 3Aiv, 3Av, 3Avi).

**Figure 3.**
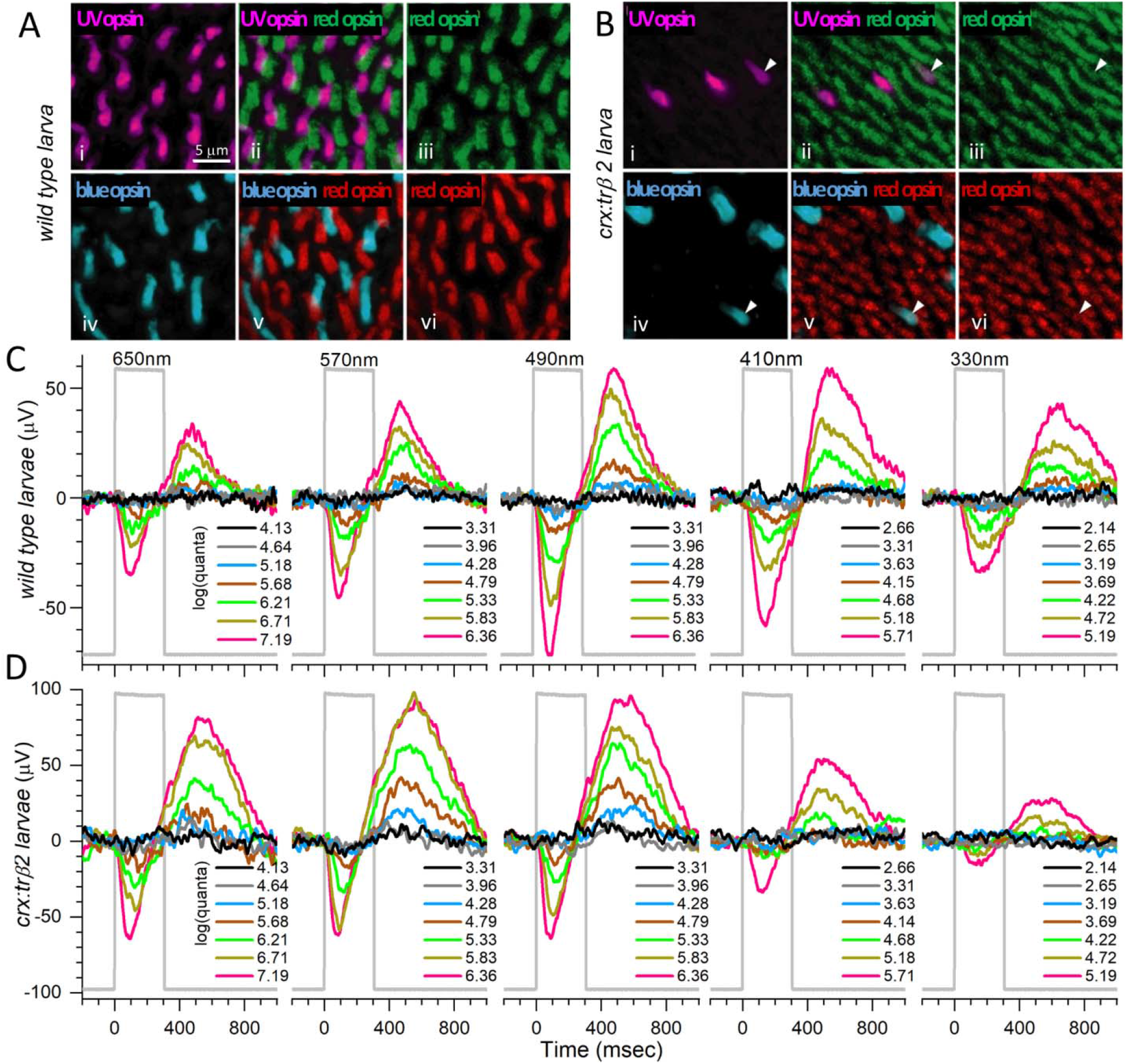
Cone distributions and spectral responses in embryonic WT and *crx:mYFP-2A-trβ2* larval eyes. ***Ai***, UV opsin (SWS1) immunoreactive cones in a WT retina. ***Aiii, Avi***, Red opsin (LWS1, LWS2) immunoreactive cones in WT retinas. ***Aiv***, Blue opsin (SWS2) immunoreactive cones in a WT retina. ***Aii***, UV and red opsins are expressed in separate cones in a WT retina. ***Av***, Red- and blue-opsins are expressed in separate cones in a WT retina. ***Bi***, UV-opsin immunoreactive cones in a *crx:trβ2* retina. ***Biii, Bvi***, Red opsin immunoreactive cones in *crx:trβ2* retinas. ***Biv***, Blue opsin immunoreactive cones in a *crx:trβ2* retina. There are fewer UV and blue cones in *crx:trβ2* retinas than in WT retinas. ***Bii***, One *crx:trβ2* cone is immunoreactive for both UV and red opsins (arrowhead). ***Bv***, Arrowhead points to a *crx:trβ2* cone immunoreactive for both red and blue opsins. ***C***, Cone signals from a WT larval eye respond to all stimulus wavelengths with largest amplitudes at 490 nm. ***D***, The larval *crx:trβ2* retina responds with maximal amplitudes at wavelengths 490, 570 and 650 nm but is less responsive than WT for 330 and 410 nm, wavelengths that stimulate blue and UV cones. ***A, B***, 5-dpf larvae. ***C, D***, 6-dpf larvae. Perfusion medium contains 20 mM Aspartate to isolate photoreceptor signals in the ERG. Five of the 9 stimulus-protocol wavelengths are illustrated. The stimulus irradiances [units of log(quanta·μm^-2^s^-1^)] appear to the right of stacked irradiance-response traces.

Red cones are denser in the retinas of the *crx:mYFP-2A-trß2* larvae (Fig. 3Biii, vi) than in WT (Fig. 3Aiii, vi). Based on figures 3A and 3B, the density of red-opsin immunoreactive cones in these 5-dpf transgenics (188,000 mm^-1^) is significantly greater than WT (72,500 mm^-1^) [*t*(10) = 18.0, *p* = 6.0 × 10^-9^]. The density of *crx:trβ2* UV opsin immunoreactive cones (7900 mm^-1^, Fig. 3Bi) is significantly less than WT (42,000 mm^-1^, Fig. 3Ai) [*t*(6) = 7.0, *p* = 0.00043)]. One of the three *crx:trβ2* UV cones (Fig. 3Bi) stains for both UV and red opsins (arrowhead, Figs. 3Bi, 3Bii, 3Biii), indicating co-expression of red and UV opsins. The density of *crx:trβ2* blue opsin immunoreactive cones (13,800 mm^-1^. Fig. 3Biv) is significantly less than WT (30,500 mm^-1^, Fig. 3Aiv) [*t*(6) = 4.7, *p* = 0.0033)]. Of four transgenic blue cones illustrated (Fig. 3Biv), one is immunoreactive for red opsin (arrowhead, Figs. 3Biv, 3Bv, 3Bvi), evidence for co-expression of red and blue opsins in a single cone. The alterations in densities of cone types resemble findings obtained in the presence of morpholino blockade of native trβ2 (Suzuki et al., 2013) suggesting the *crx:trβ2* transgene alters cone developmental patterns regardless of the activity of the native *trβ2* gene. The difference is the mixed opsin cones, which was not earlier described.

### Cone spectral signals from wild-type and crx:trβ2 retinas in larvae and adults

The spectral pattern of Aspartate-isolated cone signals from larval *crx:mYFP-2A-trß2* eyes parallels the altered cone densities. With short and UV wavelength stimulation (410 nm, 330 nm, Fig. 3C), an isolated WT eye responds with substantial signals for three of the brightest stimulus irradiances (green, yellow or red traces) but at these same wavelengths, signals from a *crx:trβ2* eye are evident at only for the two brightest irradiances (yellow and red) and are of lesser amplitude than WT (Fig. 3D). Maximal amplitudes were evoked at 490 nm in the WT eye (Fig. 3C), while amplitudes in the *crx:trβ2* eye plateaued at longer wavelengths (490, 570, and 650 nm, Fig. 3D).

The spectral differences in cone signals from adult *crx:mYFP-2A-trβ2* and WT eyecups were even more pronounced than those in larval eyes. In the WT traces, the maximal amplitude occurred at 490 nm (Fig. 4A), whereas the maximal amplitude from a *crx:trβ2* eyecup was seen at 650 nm (Fig. 4B). Amplitudes at 410 nm and 330 nm (Fig. 4B) were greatly reduced as compared to the WT control (Fig. 4A). At these short wavelengths WT signals were more sensitive than *crx:trβ2* signals, with the dimmer stimuli (gray, and blue traces, Fig. 4A, 4B) evoking deflections in WT, but only overlapping baselines in *crx:trβ2* eyecups.

**Figure 4.**
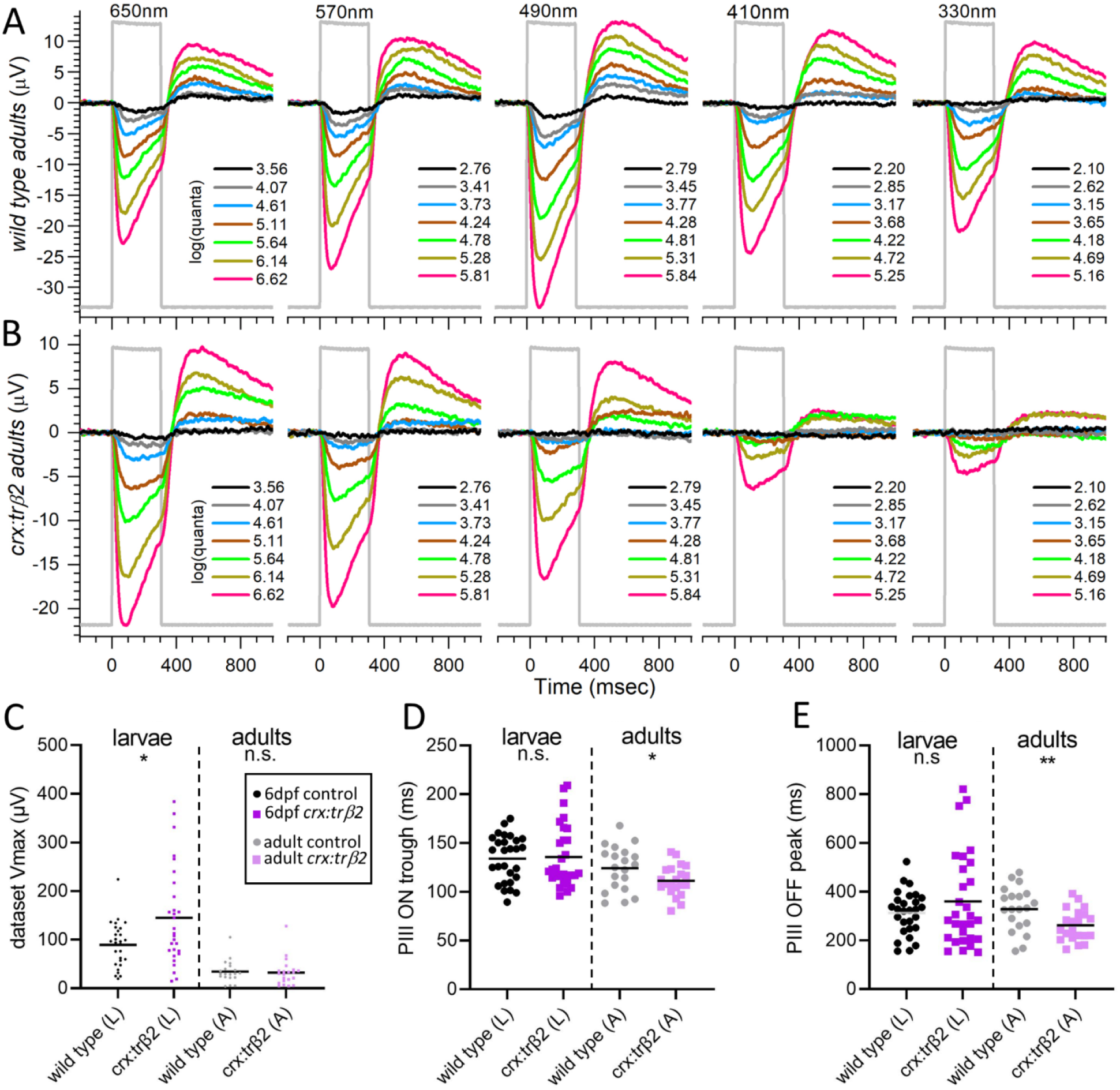
Cone spectral signals in WT and *crx:mYFP-2A-trβ2* adults. ***A***, In WT eyecups all wavelengths and most irradiances evoke signals, with a maximal response at 490 nm. ***B***, Cone signals from *crx:trβ2* eyecups are maximal at 650 nm. Response amplitudes to wavelengths that stimulate blue and UV cones (410 nm, 330 nm) are less than WT. In ***A***, and ***B***, adults are 8-18 mo. The perfusion medium contains 10 mM Aspartate to isolate photoreceptor signals. The nested responses are irradiance-response series at each wavelength, with irradiances given to the right of the traces in units of log(quanta·μm^-2^s^-1^). ***C***, ***D***, ***E***, Properties of ERG cone-PIII waveforms in larval and adult, WT and *crx:trβ2* retinas. ***C***, Distributions of maximal trough-to-peak amplitudes in larval and adult spectral datasets. ***D***, Cone-signal latency from stimulus onset to the minimum of the ON trough of the cone signal are measured on the mean responses of each 280-stimulus spectral dataset. ***E***, Cone-signal latency to the OFF peak in the mean responses of spectral datasets. OFF latencies are measured from stimulus offset. Asterisks (n.s., not significant) are probabilities that WT and *crx:trβ2* distributions differ in larvae or adults (Graphpad Prism convention, statistics given in text).

### Wild-type and crx:trβ2 waveforms in larvae and adults

The *crx:mYFP-2A-trβ2* transgene subtly changes the amplitudes and kinetics of Aspartate-isolated cone signals (cone PIII). In larvae a greater mean amplitude and greater spread of amplitudes were found (Fig. 4C). The mean of maximal peak-to-trough responses in 6 dpf spectral datasets for 10 WT larval eyes was 89.6 ±8.7 μV (28 datasets, mean and SE), while the mean of maximal amplitudes for 12 *crx:trβ2* eyes was 145.0 ±18.7 μV (29 datasets). The *crx:trβ2* amplitudes were somewhat larger [*t*(55) = 2.65,*p* = 0.0103), Fig. 4C]. The dispersion of larval amplitudes was also significantly greater for *crx:trβ2* [*F*(4.77, 28, 27) = 5.6 × 10^-5^, *p* = 0.00011]. But on reaching adulthood, *crx:trβ2* maximal PIII amplitudes became indistinguishable from WT counterparts (Fig. 4C). In 14 WT eyecups the mean of maximum peak-to-trough amplitudes was 34.4 ±5.1 μV (20 datasets). In 16 *crx:trβ2* eyecups the mean of maximal peak-to-trough amplitudes was 32.4 ±6.1 μV (21 datasets). The difference was insignificant [*t*(38) = 0.248, *p* = 0.806]. The variance in maximal amplitudes was similar [*F*(1.51, 20, 19) =0.813, *p* = 0.375]. Lesser amplitudes occur in adult eyecups than intact larval eyes as photocurrents escape around eyecup edges, whereas the larval eye is an electrically sealed system. Neither larval nor adult *crx:trβ2* amplitude distributions (Fig. 4C) show evidence of amplitude loss, a characteristic associated with retinal degenerations.

‘PIII ON trough’ is the time interval from stimulus onset to the minimum in the vitreal negative phase of the cone PIII signal. In larvae, PIII-ON-trough latencies for the *crx:mYFP-2A-trβ2* waveforms were similar to WT, but in adults the latency to the PIII ON trough was shorter (Fig. 4D). Response peak and trough timings were measured on the mean waveform for all 280 responses in a spectral dataset. In this average response, noise is minimized and interferes least with determination of extrema timing. For larval eyes the latency to the cone PIII trough for 29 *crx:trβ2* datasets (12 eyes) was 135 ±6 ms from stimulus onset (mean and SE). The trough time for 28 WT datasets (10 eyes) was 134 ±5 ms. The WT and transgenic trough times were not significantly different [*t*(55) = 0.218, *p* = 0.828] and the trough-time variances were also similar [*F*(1.80, 28, 27) = 0.935,*p* = 0.133, Fig. 4D]. In 16 adult *crx:trβ2* eyecups the mean latency to PIII ON troughs was 111 ±3 ms (21 datasets). In 14 WT eyecups the PIII trough latency was 124 ±5 ms (20 datasets). The adult *crx:trβ2* transgenics were somewhat quicker to peak [*t*(39) = 2.133,*p* = 0.039]. The variances in onset latencies was similar in *crx:trβ2* and WT adults [*F*(2.11, 19, 20) = 0.947, *p* = 0.106, Fig. 4D]. For WT controls, larval and adult trough latencies did not differ significantly [*t*(46) = 1.45, *p* = 0.155]. For *crx:trβ2* transgenics trough times for adults were significantly quicker than for larvae [*t*(48) = 3.23, *p* = 0.0022)].

‘PIII OFF peak’ (Fig. 4E) is the time interval from stimulus offset to the maximum of the upward course in the PIII OFF signal. The larval PIII-OFF-peak latencies for *crx:mYFP-2A-trβ2* (360 ±36 ms) did not differ significantly from WT controls [313 ±17 ms, *t*(55) = 1.171, *p* = 0.247], although the variance in *crx:trβ2* timing was significantly greater [*F*(4.68, 28, 27) = 0.999, *p* = 1.3 × 10^-4^]. In adults the *crx:mYFP-2A-trβ2* OFF peak (262 ±14 ms), like the ON trough, occurred significantly sooner than in WT [329 ±20 ms, *t*(39) = 2.728, *p* = 0.0095]. The variances of adult OFF-peak latencies were similar in *crx:trβ2* and WT adults [*F*(1.99, 19, 20) = 0.932, *p* = 0.135, Fig. 4E]. For WT controls, larval and adult PIII OFF peak timing was not significantly different [*t*(46) = 0.600, *p* = 0.551]. For *crx:trβ2* transgenics PIII-OFF-peak times for adults were somewhat quicker than for larvae [*t*(48) = 2.234, *p* = 0.030]. Taken together the latency distributions (Fig. 4D, 4E) show little evidence of waveform abnormalities associated with errors in cone phototransduction or retinal disease. The quicker waveforms of *crx:trβ2* adults may result from different weighting of contributing cone types.

### In crx:trβ2 larvae, red-cone signals increase but UV- and blue-cone signals diminish

Inspection of differences in larval signal amplitudes across the stimulus spectrum (Fig. 3C, 3D) suggests that the complement of cone types contributing to vitreal cone-PIII signals is altered by the *crx:mYFP-2A-trβ2* transgene. To determine the cone contributions affected, 255 models comprising all combinations of 8 cone spectral types, were fit to *crx:trβ2* and WT cumulative spectral datasets. For 6-dpf WT and *crx:trβ2* larvae, the cumulative datasets included 1858 response amplitudes (28 datasets) and 1860 amplitudes (29 datasets) respectively. Optimal models differed (WT, #111; *crx:trβ2*, #202). The residual variance of the WT model (#111) as fit to the *crx:trβ2* cumulative dataset differed from the residual variance of the *crx:trβ2* dataset fit to its own optimal model #202 [*F*(9.16,1821,1825) = 1, *p* = 0], indicating that the WT model is not an equivalently good representation of *crx:trβ2* data.

In figures 5A and 5B, amplitudes are plotted against irradiance for four of the 9 test wavelengths. Continuous curves are calculated from the optimal models for WT and *crx:mYFP-2A-trβ2* datasets. These curves are generated from a global model fit to all spectral data, not just the illustrated data. The adherence of irradiance-response curves to datapoints for individual wavelengths is an index of the ability of the global models to represent the cumulative spectral data. The distribution of points and curves is more compressed along the irradiance axis for *crx:trβ2* larvae (Fig. 5B) than for WT controls (Fig. 5A). The points and curve at 370 nm (magenta) lie to the left of the 490 nm points and curve (green) for WT spectral signals (Fig. 5A) but to the right of the 490 nm curve and points for *crx:trβ2* larvae (Fig. 5B), suggesting a loss of sensitivity for 370-nm-sensing cones (UV cones). The log of semi-saturation irradiance calculated for the R1 and R2 cones was 4.56 ±0.018 for WT larvae and 4.53 ±0.027 for *crx:trβ2* (Fig. 5A, 5B). These semi-saturation irradiances did not significantly differ [*t*(3681) = 1.858, *p* = 0.063].

**Figure 5.**
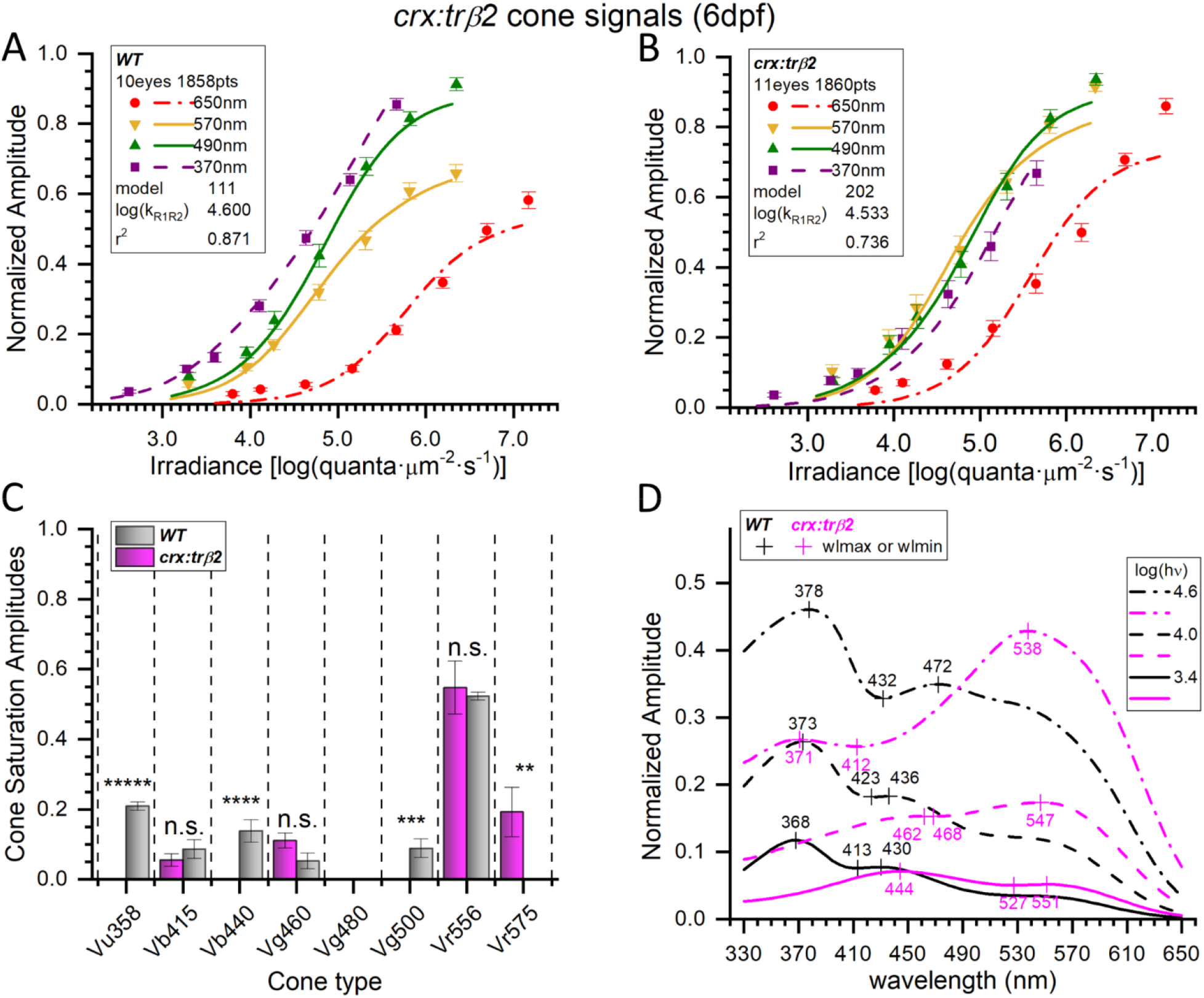
Spectral models of cumulative data from 6-dpf embryonic *crx:mYFP-2A-trβ2* and WT eyes. ***A***, WT irradiance-response datapoints, SEs, and optimal model (#111) curves as fit to 1858 spectral response amplitudes combined from 28 normalized datasets taken from 10 eyes (Fig. 1A, Eq. 1 algorithm). 370 nm, *n* = 23-28; 490-nm, *n*= 27-28; 570-nm; *n* = 23-28; 650-nm, *n* = 23-55. ***B***, *crx:trβ2* irradiance-response datapoints and optimal model (#202) curves fit to 1860 spectral points combined from 29 datasets taken from 11 eyes. Individual points: 370 nm, *n* = 22-28; 490 nm, *n* = 23-28; 570 nm, *n* = 27-28 points; 650 nm, *n* = 24-55. The *crx:trβ2* transgene moves curves and points along the irradiance axis as compared to WT. ***C***, In WT, six cone types (gray bars) were detected by the optimal model. In *crx:trβ2*, four cone signals were detected (magenta bars). Cone saturation amplitudes *[Vm_i_* values (Eq. 1, Fig.2A) ±SE] are fractions of dataset maximal amplitudes. Asterisks denote significance of differences between WT and *crx:trβ2* (one or two sample *t*-tests; n.s., not significant; GraphPad Prism convention). Vu358 (UV, one sample test): *t*(1822) = 17.8, *p* = 1.2 × 10^-65^; Vb415 (B1): *t*(3647) = 0.958, *p* = 0.338; Vb440 (B2): *t*(1822) = 4.27, *p* = 2.0 × 10^-5^; Vg460 (G1): *t*(3647) = 1.87, *p* =0.062; Vg500 (G4, one sample test): *t*(1822) = 3.33, *p* = 8.8 × 10^-4^; Vr556 (R2): *t*(3647) = 0.318, *p* = 0.751; Vr575 (R1): *t*(1825) = 2.72, *p* = 0.0067. ***D***, Spectral peaks shift to longer wavelengths for *crx:trβ2* (magenta) as compared to WT (black). Spectral curves are the modeled amplitudes that would be evoked by 3 different irradiances of constant quantal stimulation across the spectrum [3.4, 4.0, and 4.6 log(quanta·μm^-2^·s^-1^)] as spectral shapes differ with stimulus brightness. (***A***, ***B***) The log(k_R1R2_) values are modeled R1-cone and R2-cone semi-saturation irradiances in log(quanta·μm^-2^s^-1^). (***A***, ***B, C, D***) 20 mM Aspartate medium.

The inferred *Vm_i_* values for individual larval cone types (Fig. 2, Eq. 1) relate closely to functional strength within the cone-PIII signal. In figure 5C the modeled values are compared for WT and *crx:mYFP-2A-trβ2* larvae. Six cone types significantly contributed to WT cone-PIII signals (Vu358-UV, Vb415-B1, Vb440-B2, Vg460-G1, Vg500-G4, Vr556-R2), but only four cone contributions were identified in *crx:trβ2* larvae (Vb415-B1, Vg460-G1, Vr556-R2, Vr575-R1). Missing in the transgenic are UV cone signals (Vu358), B2 cone signals (Vb440), and G4 signals (Vg500). But an extra red-cone signal, R1 (Vr575), not detected in WT, became significant in *crx:trβ2* larvae, The R2 (Vr556) cone signal in this 6-dpf dataset did not differ from WT, nor did either the B1 signal (Vb415) or the G1 signal (Vg460). Overall, the modeling algorithm shows that UV-cone amplitudes were significantly reduced, red-cone amplitudes significantly increased, and among four blue and green cones signals, two were significantly reduced and two were not significantly affected.

Modeled spectral sensitivities represent the impact of not only *Vm_i_* but also *k_i_* values and include Hill-function response compressions, which flatten cone spectral functions for bright stimulation (Eq.1, Fig. 2A). This is the fullest model representation, but less interpretable in terms of individual cone contributions. The spectral curves of figure 5D are modeled response amplitudes for three levels of constant quantal stimulation across the spectrum. The strong UV signal in WT larval eyes (Fig. 5A) appears as an ultraviolet spectral peak (~370 nm) in figure 5D, regardless of stimulus brightness, despite the larger saturation amplitudes of the R2 (Vr556) cones, which, in WT controls, manifest as a long-wavelength bulge more prominent with greater constant-quantal irradiances, which better stimulate the higher semi-saturation characteristics of R2 physiology. The failure to find UV cone signals in the *crx:trβ2* larval transgenics precludes an ultraviolet spectral peak at any quantal-irradiance and allows red-cone signals to create spectral peaks (540-550 nm). The residual blue and green signals of *crx:trβ2* cooperate to create a 444 nm peak at the dimmest level of stimulation (Fig. 5D). With 255 models tested, there is not always a single best-fitting model for each dataset as judged by residual variance. For the larval WT datasets, model #95, also a 6-cone model, was indistinguishable from model #111 [*F*(0.99895, 1851, 1851) = 0.491, *p* = 0.982]. This model replaces the low amplitude G4 (Vg500) cone amplitude with a similar amplitude G3 (Vg480) signal. The WT five-cone model #103 was also indistinguishable [*F*(0.99719, 1851, 1852) = 0.476, *p* = 0.952]. This model omits the low-amplitude G1 (Vg460) cone signal. For the *crx:trβ2* datasets, model #201 was indistinguishable from model #202 [*F*(0.999, 1856,1856) = 0.491, *p* = 0.9822]. This model substitutes a low-amplitude UV-cone signal (Vu358) for the low-amplitude B1-cone signal (Vb415).

### Adult crx:trβ2 zebrafish are red cone dichromats

Only the red R1 (Vr575) and R2 (Vr556) cone signals made significant contributions to the adult cone-PIII responses of *crx:mYFP-2A-trβ2* (Fig. 6C). A five-cone WT model including blue-, green- and red-cones best represented adult controls (Vb415-B1, Vg460-G1, Vg500-G4, Vr556-R2, Vr575-R1). The cumulative datasets included 14 eyecups from WT adults and 16 from *crx:trβ2* adults (WT, 20 datasets, 1400 responses; *crx:trβ2*, 21 datasets, 1470 responses). When the adult WT model (#234) is fit to the adult *crx:trβ2* dataset, the residual variance is larger than the variance for the optimal *crx:trβ2* model (#192) [*F*(1.923,1443,1445) = 1, *p* = 0] indicating different spectral patterns are operating. The *crx:trβ2* transgene alters the positioning of adult irradiance-response data points and model curves (Figs. 6A, 6B). The 370 and 490 nm curves and points (magenta and green) lie to the left side of the 570 nm curve and points (yellow) in WT adults but shift to the right side of the 570 nm curve and points in *crx:trβ2* adults, a diminution of UV- and mid-wavelength sensitivity. The calculated log of semi-saturation irradiance for long-wavelength-sensitive cones (R1, R2) was 4.49 ±0.033 for WT adults and 4.69 ±0.022 for *crx:trβ2* adults (Fig. 6A, 6B), *crx:trβ2* being significantly less sensitive [*t*(2818) = 5.17, *p* = 2.5 × 10^-7^]. The modeled amplitudes of cones with significant signals appear in Fig. 6C. In the adult WT model, significant blue- and green-cone signals (B1, Vb415 and G1, Vg460) are detected, but these signals are not significant in adult *crx:trβ2* transgenics, where there are only red-cone signals (R2, Vr556 and R1 Vr575). In *crx:trβ2*, R1 cones contribute a significantly larger amplitude signal than R2 cones [180%, *t*(2818) = 3.92, *p* = 0.000092)], whereas in the WT controls, R1 and R2 signal amplitudes were not significantly different.

**Figure 6.**
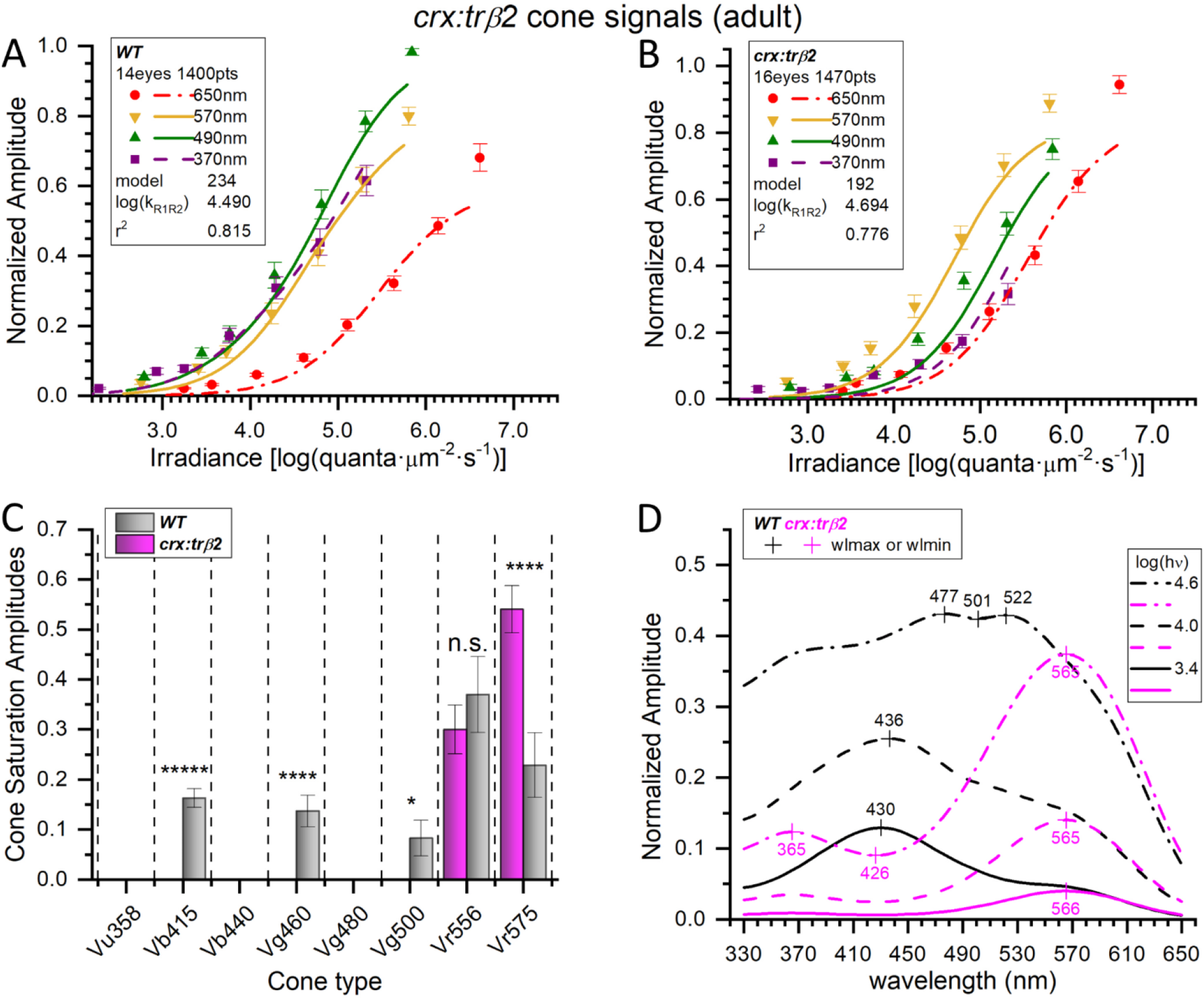
*crx:mYFP-2A-trβ2* adults are red-cone dichromats. ***A***, WT cone-PIII irradiance-response curves and datapoints at four stimulus wavelengths. The 370-, 490-, 570, and 650-nm amplitudes are means (±SE) from the cumulative dataset (*n* = 20 except at 650-nm where *n* = 20 or 40). Curves are generated from the optimal model (#234) fit to a cumulative 1400 responses at 7 irradiances for each of 9 wavelengths as compiled from 20 spectral datasets acquired in 14 WT adult eyecups. ***B***, The *crx:trβ2* gain-of-function transgene moves irradiance-response points and model curves along the irradiance axis as compared to WT. The model generating the curves (#192) is fit to 1470 cumulative responses compiled from 21 datasets acquired from 16 *crx:trβ2* eyecups. The amplitudes (±SE) are means (370-, 490-, and 570, 650 nm, *n* = 21 except for 650 nm, *n* = 21 or 42). ***C***, Signals from five blue, green, or red cone types were detected in WT adults (gray bars) but signals from only the two red-cone types were detected in *crx:trβ2* eyecups (magenta bars). Fit values of cone saturation amplitudes (*Vm_i_*, Eq. 1, Fig. 2A) are plotted on a dataset-normalized scale (±SE). Except for Vr556 (LWS2) cone types differed significantly in amplitudes between WT and *crx:trβ2* (asterisks, GraphPad Prism convention, n.s., not significant). Vb415 (B1, one sample test): *t*(1373) = 8.77, *p* = 5.4 × 10^-18^; Vg460 (G1, one sample test): *t*(1373) = 4.36, *p* = 1.4 × 10^-5^; Vg500 (G3, one sample test): *t*(1373) = 2.30, *p* = 0.021; Vr556(R2): *t*(2818) = 0.782, *p* = 0.434; Vr575 (R1): *t*(2818) = 3.91, *p* = 9.2 × 10^-5^. ***D***, Model spectral curves for adult WT (black) and *crx:trβ2* (magenta) eyecups. The *crx:trβ2* transgene shifts sensitivity peaks to long wavelengths at all stimulus irradiances. Curves are modeled for constant quantal stimuli at 3.4, 4.0, and 4.6 log(quanta ·μm^-2^·s^-1^). (***A, B***) The log(k_R1R2_) values are the irradiance semi-saturation values in log(quanta·μm^-2^s^-1^) for both R1 and R2 cones. (***A, B***, ***C, D***). 8-18-month adults, 10 mM Aspartate medium.

The modeled spectral curves for the *crx:mYFP-2A-trβ2* adult red-cone dichromat (Fig. 6D) show long-wavelength-peaking functions at 565 nm at all quantal stimulation levels, while WT spectral peaks occur between 430 nm and 522 nm, depending on stimulus brightness. Except at wavelengths greater than 570 nm, WT control amplitudes are greater than transgenic, due both to the greater sensitivity of WT cones overall, and to the presence of mid- and short-wavelength (blue and green) cones, absent in the transgenic. As judged by residual variance no other models were indistinguishable from the illustrated ones for either *crx:trβ2* or WT adults (*F*-tests for all other models, *p* < 0.95).

### Larval gnat2:trβ2 cone types and spectral signals

The *gnat2:mYFP-2A-trβ2;mpv17* transgenic was developed to test whether red cones could be restored to a population of cones already differentiated under morpholino blockade of the native *trβ2* gene, which prevented red-cone development (Suzuki et al., 2013). The *gnat2* locus codes for cone transducin α, a gene product only expressed in differentiated and functional cone cells. It is expressed by 4 dpf. Suzuki et al (2013) found the *gnat2:trβ2* transgene induced a supra-normal 5-dpf density of red-opsin immunoreactive cones in the absence of native trβ2, but unlike the *crx:mYFP-2A-trβ2* rescue, green-(Rh2), blue- (SWS2) and UV- (SWS1) opsin immunoreactive cone densities were normal (Suzuki et al., 2013).

A similar pattern of larval opsin immunoreactivity is seen without morpholino blockade of native trβ2. In the control retina (*roy orbison mpv17^-/-^*) both UV and red cones (Fig. 7Ai, 7Aiii) stain with opsin antibodies. Superposition of red- and UV-cone mosaics (Fig.7Aii) shows no red- and UV- opsin overlap, with immunoreactivity for each opsin expressed in a separate cone cell. The control larval blue cones also exist in a mosaic pattern separate from red cones (Figs. 7Aiv, 7Av, 7Avi). The *gnat2:trβ2* retina shows dense representations of UV- and blue-opsin-immunoreactive cones (Figs. 7Bi, 7Biv). Red-opsin immunoreactive cones appear denser in *gnat2:trβ2* than in *mpv17^-/-^* control retinas (Figs. 7Biii, 7Bvi). The 12-dpf *gnat2:trβ2* red cone density (95,500 mm^-1^) was significantly greater than the control strain (51,400 mm^-1^) [*t*(14) = 6.7, *p* = 0.000011, Figs. 7Aiii, 7Avi, 7Biii, 7Bvi]. The densities of UV- and blue-opsin immunoreactive cones were similar in *gnat2:trβ2* and controls. The *gnat2:trβ2* UV-cone density (34,252 mm^-1^) was not significantly different from *mpv17^-/-^* (23,435 mm^-1^) [*t*(6) = 1.7, *p* = 0.143], nor was the *gnat2:trβ2* blue-cone density (21,633 mm^-1^) different from *mpv17^-/-^* (27,041 mm^-1^) [*t*(6) = 1.6, *p* = 0.168]. In these 12-dpf *gnat2:trβ2* transgenics (Fig. 7B), UV- and blue-opsin immunoreactive cones were frequently immunoreactive for red opsins (arrowheads), with 2/3 of cones being mixed opsin types.

**Figure 7.**
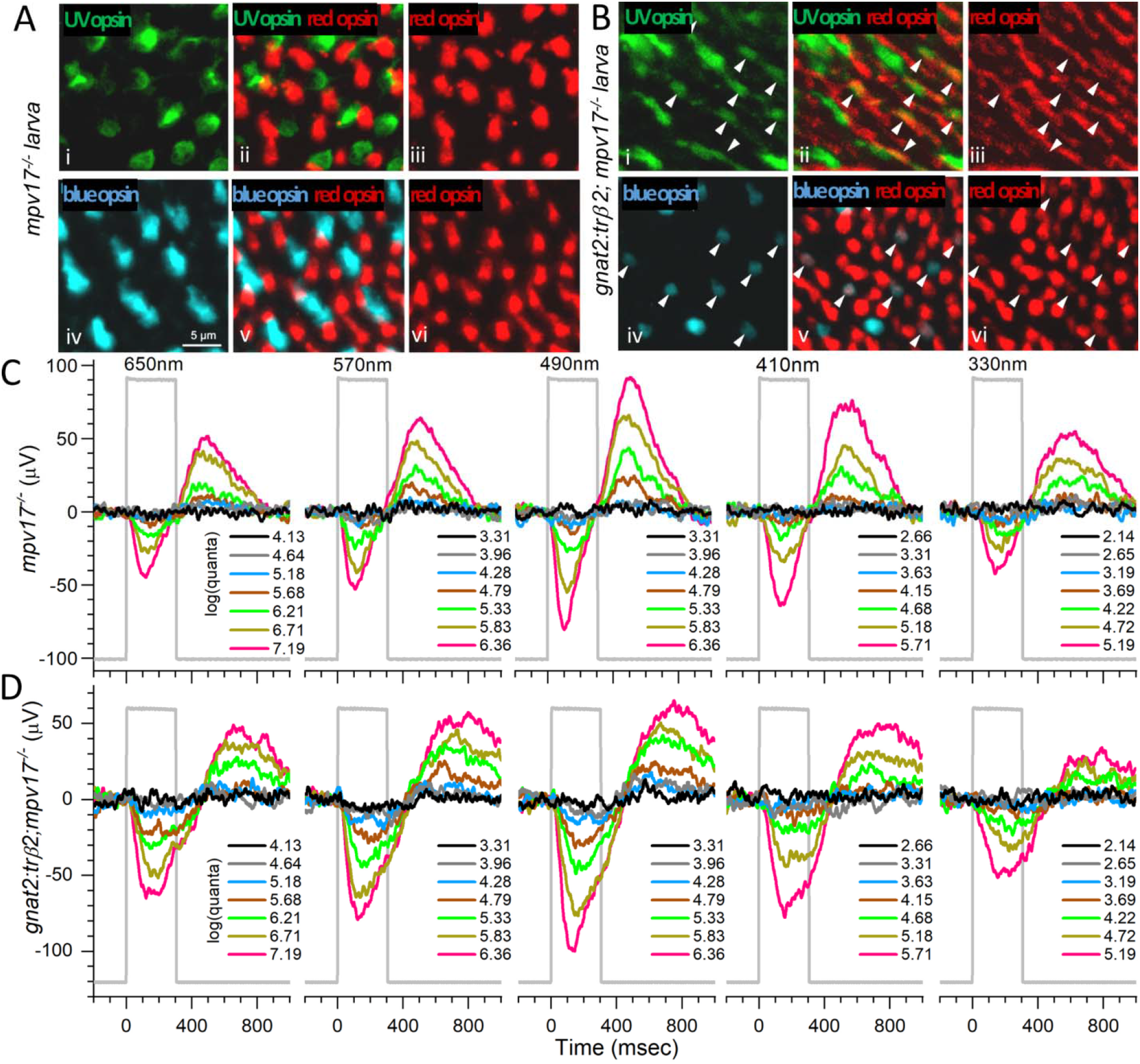
Larval cone distributions and spectral responses in *gnat2:mYFP-2A-trβ2;mpv17^-^* eyes and controls. ***Ai***, UV opsin (SWS1) opsin immunoreactive cones in control (*roy orbison mpv17^-/-^*) retinas. ***Aiv***, Blue opsin (SWS2) immunoreactive cones in control retina. ***Aiii***, ***Avi***, Red (LWS1, LWS2) opsin immunoreactive cones in control retinas. ***Aii***, UV and red opsins are expressed in separate cones in control retina. ***Av***, Blue and red opsins are expressed in separate cones in control retina. ***Bi***, UV opsin immunoreactive cones in *gnat2:trβ2* retina. ***Biv***, Blue opsin immunoreactive cones in *gnat2:trβ2* retina. ***Biii***, ***Bvi***, Red opsin immunoreactive cones in *gnat2:trβ2* retinas. ***Bii***, ***Bv***, Red opsins are expressed in the blue or UV opsin immunoreactive cones of *gnat2:trβ2* retinas (arrowheads, ***Bii***, ***Bv***). ***A***, ***B***, 12-dpf larvae. ***C***, ***D***, Cone signals in *mpv17^-/-^* control and *gnat2:trβ2* retinas are robust at all wavelengths, with greatest amplitudes at 490 nm. The perfusion medium contains 20 mM Aspartate to isolate cone signals. Stimulus irradiances [log(quanta·μm^-2^s^-1^)] appear in the legends to the right of irradiance-response trace stacks. 6-dpf larvae.

Densities of cone types and mixed opsin patterns were similar earlier in development. At 7 dpf, the density of red-opsin immunoreactive cones in the was 70,800 mm^-1^, significantly greater than the *mpv17^-/-^*control (52,300 mm^-1^) [*t*(10) = 13.0, *p* = 1.3 × 10^-7^]. The densities of blue-opsin immunoreactive cones were similar [*gnat2:trβ2*, 22,000 mm^-1^; *mpv17^-/-^* 24,900 mm^-1^, *t*(7) = 1.9, *p* = 0.096], but the density of 7-dpf *gnat2:trβ2* UV immunoreactive cones was slightly less [*gnat2:trβ2*, 36,900 mm^-1^; *mpv17^-/-^*, 46,900 mm^-1^, *t*(8) = 3.4, *p* = 0.010]. Mixed blue-red and UV-red opsin expression was noted in about 10% of 7-dpf blue- or UV-opsin immunoreactive cones.

### Larval and adult cone spectral signals from control and gnat2:trβ2 retinas

Despite greater red-cone density and numerous mixed opsin cones (Fig. 7B), the spectral patterns of larval, Aspartate-isolated cone signals (cone-PIII) from in-vitro *gnat2:mYFP-2A-trβ2;mpv17^-/-^* eyes are similar to the *mpv17^-/-^* controls. In the trace recordings of figure 7C, an *mpv17^-/-^* control larval eye produces substantial signals at all wavelengths, with maximal amplitudes for the 490 nm stimulus. A *gnat2:trβ2* larval eye gives a similar amplitude pattern for the test wavelengths and irradiances (Fig. 7D), with maximal amplitudes at 490 nm.

In an adult *mpv17^-/-^* control eyecup (Fig. 8A), the spectral pattern of cone-PIII ERG waveforms resembles those of larvae, with a broad range of responsiveness across wavelengths, and the 490-nm stimuli yielding an amplitude peak. An adult *gnat2:mYFP-2A-trβ2;mpv17^-/-^* eyecup also shows a broadly responsive spectral pattern (Fig. 8B), but greater amplitudes are evoked at long wavelengths (570 nm, 650 nm), suggesting that the underlying input signals from red, green, blue and UV cone types have undergone developmental transformation.

**Figure 8.**
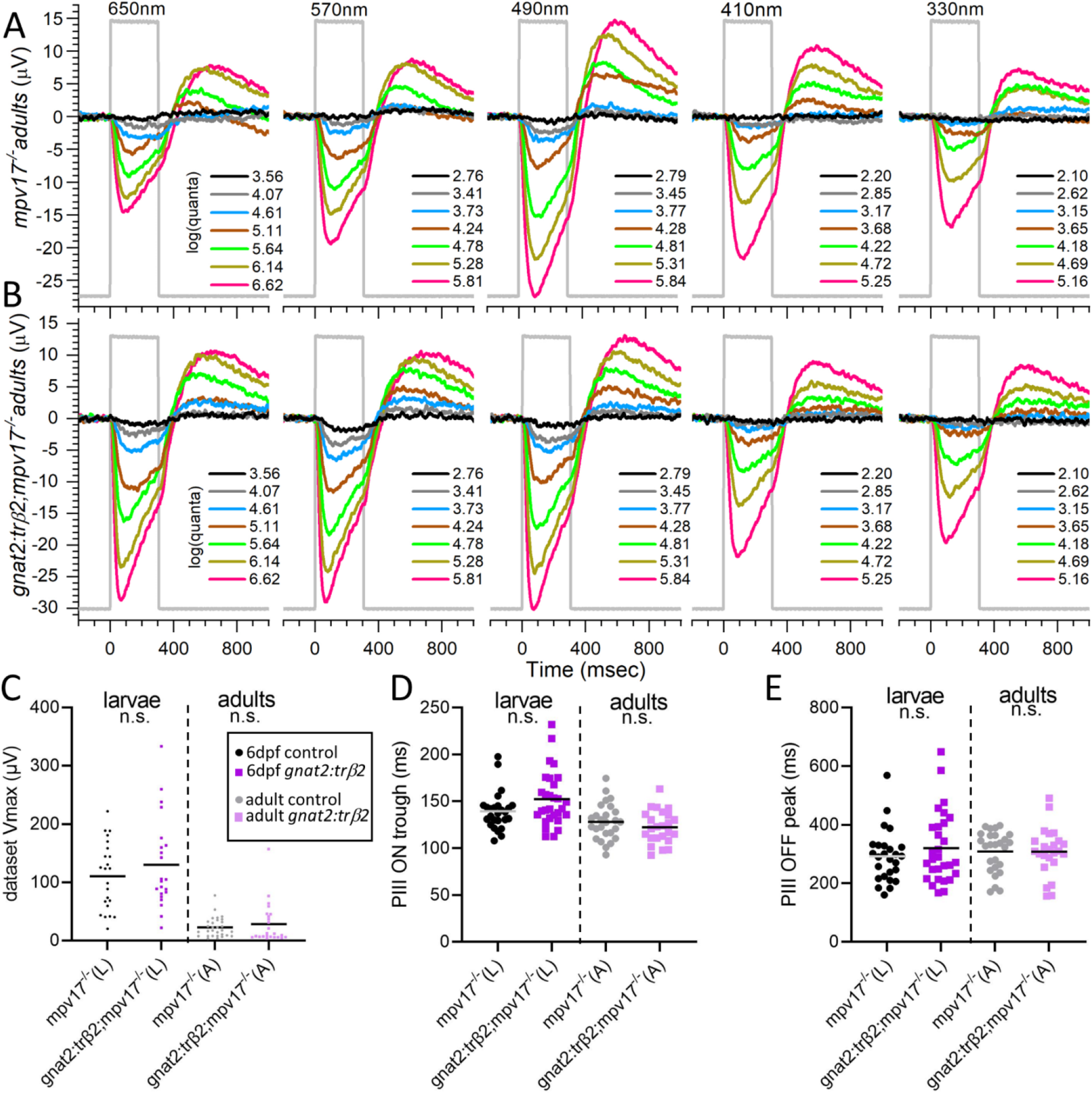
Cone PIII spectral properties in control and *gnat2:mYFP-2A-trβ2;mpv17^-/-^* adults. ***A***, Cone signals in an *mpv17^-/-^* control eyecup are evoked by all illustrated stimulus wavelengths and all but the dimmest irradiances (black). The largest amplitudes occur at 490 nm. ***B***, Cone signals from a *gnat2:trβ2* eyecup are stimulated by all wavelengths and irradiances. Greatest amplitudes occur at 650, 570, and 490 nm. ***A***, ***B***, Adults are 8-18 mo. Eyecups are perfused with medium containing 10 mM Aspartate to isolate retinal cone signals. Stimulus irradiances [log(quanta·um^-2^s^-1^)] appear in the legends to the right of irradiance-response waveform stacks. ***C***, Voltage distributions of maximal trough-to-peak amplitudes found in spectral datasets. ***D***, Cone-signal latencies from stimulus onset to the minimum in the ON trough. ***E***, Cone-signal latency from stimulus offset to the OFF peak. ***C***, ***D***, ***E***, The ‘n.s.’ labels on all distributions indicate that that *mpv17^-/-^* controls did not significantly differ from *gnat2:trβ2* transgenics in waveform characteristics (*t*-test and *p*-values given in text). Peak amplitudes (dataset Vmax), trough (PIII ON trough) and peak (PIII OFF peak) latencies are measured on the mean waveforms from each 70-stimulus spectral dataset.

### Similar waveforms from control and gnat2:trβ2 larvae and adults

There was little difference in the distribution of amplitudes or waveform kinetics between *gnat2:mYFP-2A-trβ2;mpv17^-/-^* transgenic and the *mpv17^-/-^* control strain (Figs. 8C, 8D, 8E). In both larvae and adults, the maximal ERG PIII amplitudes were similar for the *gnat2:trβ2* transgenics and controls. The 6-dpf larval mean of dataset-maximum peak-to-trough responses (23 datasets, 15 eyes) for *mpv17^-/-^* controls was 110.5 ±12.1 μV (mean and SE). For *gnat2:trβ2* larvae, the mean of dataset-maximum peak-to-trough amplitudes was 130.3 ±16.5 μV (22 datasets, 17 eyes). Maximal amplitudes of the larval *gnat2:trβ2* signals were not significantly different from those of the background strain [*t*(43) = 0.969, *p* = 0.338)]. The variances in maximal dataset amplitudes were similar in transgenic and control larvae [*F*(1.79, 21, 22) = 0.908, *p* = 0.185].

Adult *gnat2:mYFP-2A-trβ2;mpv17^-/-^* maximal cone PIII amplitudes were not distinguishable from controls (Fig. 8C). In 16 *mpv17^-/-^* adult eyecups the mean of dataset-maximum peak-to-trough amplitudes was 22.5 ±3.1 μV (30 datasets). In 15 *gnat2:trβ2* eyecups the mean of dataset-maximum peak-to-trough amplitudes was 28.2 ±7.2 μV (24 datasets). The difference was not significant [*t*(52) = 0.780, *p* = 0.439]. The variance in adult transgenic peak amplitudes was greater [*F*(4.38, 23, 29) = 0.9999, *p* = 0.00025].

Examining kinetics of the larval and adult dataset mean waveforms, the latency to the cone-PIII trough minimum and the variance in its timing differed little between 6-dpf larval *gnat2:trβ2* eyes and the *mpv17^-/-^* controls (Fig. 8D). The onset time for the larval cone-PIII trough for *gnat2:trβ2* eyes (29 datasets, 17 eyes) was 152 ±5 ms from stimulus onset (mean and SE). The cone-PIII trough latency for larval eyes from the *mpv17^-/-^* controls (26 datasets, 16 eyes) was 139 ±4 ms. The trend, though not significant, for larval *gnat2:trβ2* onset kinetics was towards greater delay in the PIII ON trough [*t*(53) = 1.89,*p* = 0.065)]. Larval *gnat2:trβ2* eyes trended, though not significantly, towards greater variability in trough latencies [*F*(2.09, 28, 25) = 0.9669, *p* = 0.067].

For adults the mean time to the PIII ON trough was 122 ±4 ms in *gnat2:trβ2* eyecups (24 datasets, 15 eyes) and 128 ±4 ms in the *mpv17^-/-^* controls (26 datasets, 16 eyes). The differences were not significant [*t*(48) = 1.11,*p* = 0.274)]. The variances in adult trough latencies for *gnat2:trβ2* and background strain were similar [*F*(1.23, 25, 23) = 0.689, *p* = 0.623]. Adult trough latencies were significantly quicker than larvae for both *mpv17^-/-^* controls and *gnat2:trβ2* transgenics *[mpv17^-/-^: t*(50) = 2.03,*p* = 0.047; *gnat2:trβ2 t*(51) = 4.39, *p* = 0.000057].

The mean cone-PIII OFF peak timing and variance in larval and adult *gnat2:mYFP-2A-trβ2 mpv17^-/-^* did not differ from controls (Fig. 8E). In the mean waveforms for the 6-dpf *gnat2:trβ2* larval datasets peaked at 321 ±22 ms after stimulus offset (29 datasets, 15 eyes). For the larval *mpv17^-/-^* controls, the OFF peak occurred at 292 ±18 ms (26 datasets, 16 eyes). These latencies were not significantly different [*t*(53) = 0.980, *p* = 0.332]. The variances in larval OFF peak timing for transgenic and control were similar [*F*(1.83, 28, 25) = 0.935,*p* = 0.132]. For adult datasets, the OFF peak of the *gnat2:trβ2* mean waveforms gave delays of 308 ±17 ms from stimulus offset (24 datasets, 15 eyes). In the adult *mpv17^-/-^* control datasets, the delay was 309 ±14 ms. The *gnat2:trβ2* adult OFF-peak timing did not differ from controls [*t*(48) = 0.031, *p* = 0.976)]. The variances in adult OFF peak timings were not different [*F*(1.34, 23, 25) = 0.763, *p* = 0.473]. Adult OFF-peak latencies did not differ significantly from larvae for either *mpv17^-/-^*controls or *gnat2:trβ2* transgenics *[mpv17^-/-^*: *t*(50) = 0.75,*p* = 0.459; *gnat2:trβ2: t*(51) = 0.43,*p* = 0.670]. Overall, the *gnat2:trβ2* transgene has little effect on cone-PIII amplitudes or waveforms. There is no electrical evidence of degenerative disease.

### In gnat2:trβ2 larvae the transgene increases R2-cone signals

The larval cone PIII spectral responses of *gnat2:mYFP-2A-trβ2;mpv17^-/-^* are similar to the *mpv17^-/-^* control strain (Fig. 7C, 7D). Signal loss in the UV and signal gain at long wavelengths, as seen *crx:mYFP-2A-trβ2* larvae (Figs. 3C, 3D), are not as evident. Modeling suggests subtle changes. The optimal model for *gnat2:trβ2* is #79, and optimal control model is #77. Applying the control model #77 to the *gnat2:trβ2* dataset gives a greater residual variance than model #79 [*F*(1.21, 1506, 1511) = 0.999,*p* =.00021], indicating that cone inputs generating the *gnat2:trβ2* cumulative dataset differ from control. In figures 9A and 9B, amplitudes are plotted against irradiance for four of the 9 test wavelengths and compared to continuous curves calculated from best-fitting models. These are fit to 1945 points (15 eyes, 28 spectral datasets) for *mpv17^-/-^* controls and 1540 points (17 eyes, 22 spectral datasets) for *gnat2:trβ2*.The distribution of points and curves at 570, 490 and 370 nm are more overlapping along the irradiance axis for *gnat2:trβ2* larvae than are the same wavelength points and curves for the control strain. Points and curves at 370 and 490 nm (magenta, green) lay to the left of the 570 nm points and curve (yellow) for *mpv17^-/-^* controls but coincide with the 570-nm curve for *gnat2:trβ2*, suggesting less sensitivity from short- and mid-wavelength-sensitive cones. The logs of semi-saturation irradiances for the red (R1, R2) cones were 4.50 ±0.02 for *mpv17^-/-^* larvae and 4.51 ±0.02 for *gnat2:mYFP-2A-trβ2;mpv17^-/-^* (Fig. 9A, 9B), nearly identical [*t*(3422) = 0.251, *p* = 0.802)].

**Figure 9.**
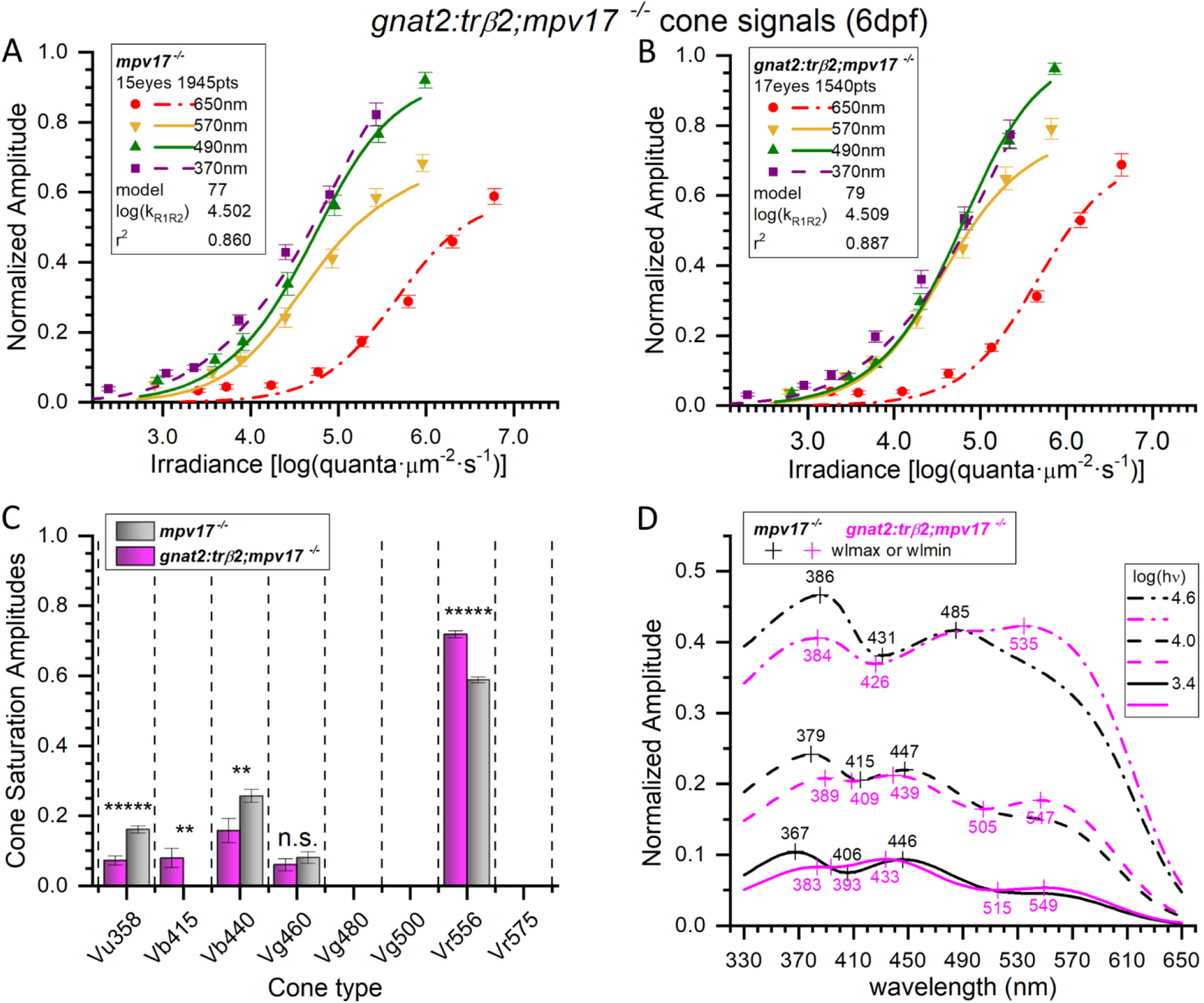
Spectral properties of embryonic cone signals from *gnat2:mYFP-2A-trβ2;mpv17^-/-^’* larval eyes and *mpv17^-/-^* controls. ***A***, Control (*mpv17^-/-^*) irradiance-response points and optimal model curves. The WT model (#77) was fit to 1945 spectral response amplitudes, combined from 28 datasets taken from 15 eyes. The points are subsets of the cumulative data. The 370-, 490- and 570-nm amplitudes are means (±SE), *n* = 27 or 28; the 650-nm means, *n* = 28 or 56. ***B***, The *gnat2:trβ2* irradiance-response points and model curves are more bunched together than the 370 nm, 490 nm, and 570 nm irradiance-response points and curves. The *gnat2:trβ2* model (#79) was fit to 1540 spectral points, combined from 22 datasets taken from 17 eyes. The 370-nm, 490-nm, and 570-nm amplitudes are means (±SE), *n* = 22; 650-nm, *n* = 22 or 44. ***C***, Four cone types were significant in cone-signals isolated from *mpv17^-/-^* control eyes (gray bars) but five types proved significant in *gnat2:trβ2* eyes (magenta bars). Cone saturation amplitudes (*Vm_i_* Eq. 1, Fig. 2A) are fractions of maximal dataset amplitudes (±SE). Asterisks represent the significance of amplitude differences between *mpv17^-/-^* control and *gnat2:trβ2* (GraphPad Prism convention, n.s., not significant). Vu358 (UV): *t*(3422) = 5.37, *p* = 8.7 × 10^-8^; Vb415 (B1, one sample test): *t*(1511) = 2.90, *p* =0.0038; Vb440 (B2): *t*(3422) = 2.62, *p* = 0.0081; Vg460 (G1): *t*(3422) = 0.851, *p* = 0.395; Vr556 (R2): *t*(3422) = 9.94, *p* = 5.6 × 10^-23^. ***D***, Model spectral curves for *mpv17* controls and *gnat2:trβ2* larval eyes for constant quantal irradiances of 3.4, 4.0, and 4.6 log(quanta·μm^-2^s^-1^). There is greater long-wavelength sensitivity, and lesser short-wavelength sensitivity in the trβ2 gain-of-function transgenic. ***A***, ***B***, The log(k_R1R2_) values are the modeled R1- or R2-cone semi-saturation irradiances in log(quanta·μm^-2^s^-1^). ***A***, ***B***, ***C***, ***D***, 6-dpf larvae, 20 mM Aspartate medium.

Four cone signals are identified in larval *mpv17^-/-^* controls, but five were found in *gnat2:mYFP-2A-trβ2* siblings (Fig. 9C). Significantly reduced in *gnat2:trβ2* are UV (Vu358) and B2 (Vb440) cone amplitudes. Significantly increased are B1 (Vb415) and R2 (Vr556) cone signals. The G1 cone signal (Vg460) was not significantly affected. The larval *mpv17^-/-^* control and *gnat2:trβ2* model-generated spectral sensitivities are similar (Fig. 9D), but with increased long-wavelength and decreased short-wavelength sensitivities in *gnat2:trβ2*. This leads to long-wavelength spectral peaks between 535 nm and 549 nm for *gnat2:trβ2*, not seen in the *mpv17^-/-^* controls. While there is a definite spectral shift towards long-wavelength sensitivity in *gnat2:trβ2* larvae (Fig. 9D), the extent is subtle compared to *crx:trβ2* larvae (Fig. 5D). Of the 255 models fit to the 6-dpf larval *gnat2:trβ2* combined dataset none were deemed to fit equally as well as model #79 based on residual variance, that is by an *F*-test with *p ≥* 0.95. Two were indistinguishable from the best-fit model 77 for the *mpv17^-/-^* controls. Model 103 [*F*(0.9992, 1939, 1940) = 0.493, *p* = 0.986], employed an additional 5^th^ cone substituting G4 for G1 and adding B1. Model 101 [*F*(0.9972, 1940, 1940) = 0.475, *p* = 0.95] substituted G4 for G1. All three indistinguishable control models agreed on the presence of UV, B2 and R2 cone signals in *mpv17^-/-^* controls.

### Adult gnat2:trβ2 retinas generate large R1 red-cone signals

In the adult *gnat2:mYFP-2A-trβ2;mpv17^-/-^* eyecup (Fig. 8B), large-amplitude responses to 570- and 650-nm stimuli suggest an increased contribution of red-cones as compared to either *gnat2:trβ2* larvae (Fig. 7B) or adult controls (Fig. 8A). To determine the cone-type composition of adult *gnat2:trβ2* retinal cone signals we searched for the best models to represent the *gnat2:trβ2* cumulative spectral dataset. Trough-to-peak amplitudes for all eyes and datasets (*mpv17^-/-^*, 13 eyes, 23 datasets, 1610 responses; *gnat2:trβ2*,11 eyes, 20 datasets, 1400 responses) were fit to the 255 combinations of 8 cones. Model #219 fit best for *mpv17^-/-^* control eyecups, and model #202, for *gnat2:trβ2*. When the adult *gnat2:trβ2* cumulative dataset was fit to the control model #202, the residual variance was significantly greater than with the optimal *gnat2:trβ2* model, indicating the spectral properties of the *gnat2:trβ2* gain-of-function dataset differed significantly from the control [*F*(1.942, 1370, 1374) = 1, *p* = 0].

For both points and optimal model curves the *gnat2:trβ2* transgene alters adult irradiance-response characteristics (Fig. 10). In the control dataset (*mpv17^-/-^*), the 570 nm points and curve (yellow) show less sensitivity than either the 370 or 490 nm points and curves (magenta, green), but greater sensitivity in the *gnat2:trβ2* (Figs. 10A, 10B). In *gnat2:trβ2*, brighter stimuli are required to elicit responses in the UV (370 nm) and mid spectrum (490 nm). The fit of log of semi-saturation irradiances for the red cones (R1, R2) were 4.30 ±0.026 for adult controls and 4.43 ±0.026 for *gnat2:trβ2* adults (Figs. 10A, 10B), a significantly greater sensitivity (~35%) than for *gnat2:trβ2* adults [*t*(2953) = 3.44,*p* = 0.0006)].

**Figure 10.**
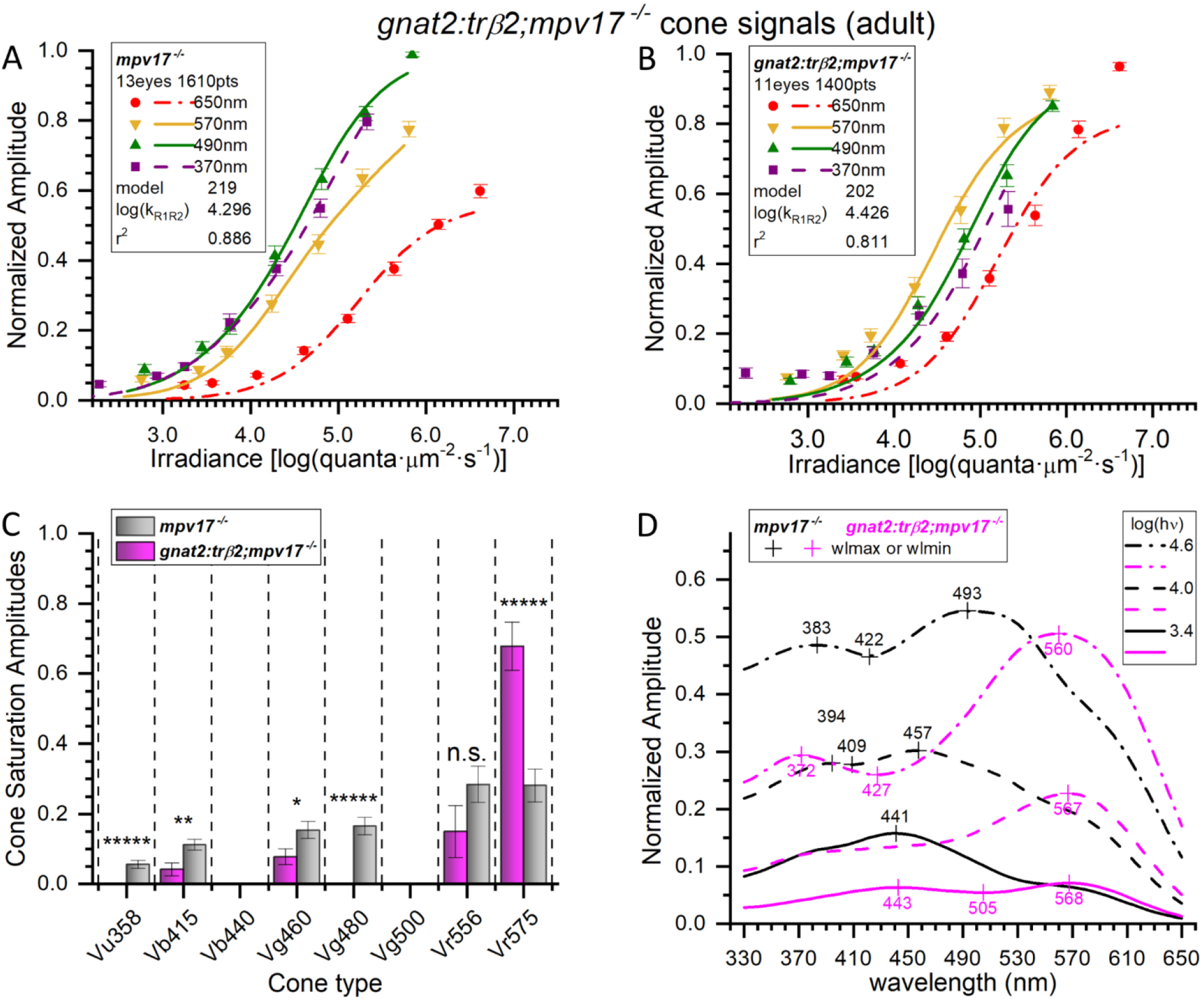
Adult spectral signals from *gnat2:mYFP-2A-trβ2;mpv17^-/-^* transgenics and *mpv17^-/-^* controls. ***A***, Control (*mpv17^-/-^*)irradiance-response amplitudes and model curves at four wavelengths. The optimal model (#219) was fit to 1610 responses (all wavelengths) combined from 23 datasets taken from 13 eyecups. The 370-nm, 490-nm and 570-nm amplitudes are means (±SE), *n*= 23; 650-nm, *n* = 23 or 46. ***B***, Irradiance-response amplitudes, and model curves, for *gnat2:trβ2*. The transgene shifts 370-nm and 490-nm irradiance-response functions from the left of the 570-nm curve (control) to the right of the 570-nm curve (transgenic). The spectral algorithm was fit to 1400 points combined from 20 datasets accumulated from 11 eyecups. The 370-nm, 490-nm, and 570-nm amplitudes are means (±SE), *n* = 23; 650-nm, *n* = 23 or 46. ***C***, In adults, significant signals from six cone types were detected in the cone responses of *mpv17^-/-^* control eyecups (gray bars). Four were found in *gnat2:trβ2* transgenics (magenta bars). Cone saturation amplitudes (*Vm_i_* Eq. 1, Fig.2A) are fractions of dataset maximum amplitudes (±SE). Asterisks (or n.s., not significant) represent the significance of differences (GraphPad Prism convention). Vu358 (UV, one sample test): *t*(1579) = 4.81, *p* = 1.7 × 10^-6^; Vb415 (B1): *t*(2983) = 2.97, *p* = 0.0030; Vg460 (G1): *t*(2983) = 2.26, *p* = 0.024; Vg480 (G3, one sample test): *t*(1579) = 6.59, *p* = 6.1 × 10^-11^; Vr556 (R2): *t*(2983) = 1.52, *p* = 0.127; Vr575 (R1): *t*(2983) = 4.87, *p* = 1.2 × 10^-6^. ***D***, Adult spectral curves for *mpv17^-/-^* and *gnat2:trβ2* cone signals. The transgene shifts sensitivity peaks to long wavelengths for all constant-quantal irradiances [3.4, 4.0, and 4.6 log(quanta·μm^-2^s^-1^)]. ***A***, ***B***, The log(k_R1R2_) values are R1 or R2 cone semi-saturation irradiances in log(quanta μm^-2^s^-1^). ***A***, ***B***, ***C***, ***D***, 8-18-month adults, 10 mM Aspartate medium.

In the best-fitting model for *gnat2:mYFP-2A-trβ2;mpv17^-/-^* adults, the long-wavelength R1 cone (Vr575) contributes the largest amplitude signal with the R2 amplitude (Vr556) being only 22% as large, significantly less than R1 [*t*(2748) = 5.23, *p* = 2.6 × 10^-17^)], and R1 amplitudes are only marginally significant [*t*(1375) = 2.02,*p* = 0.044), Fig. 10C]. In the *mpv17^-/-^* control, both R1 and R2 red cone amplitudes were highly significant [Vr575-R1, *t*(1580) = 6.01, *p* = 4.1 x 10^-18^; Vr556-R2, *t*(1580) = 5.52, *p* = 8.0 x 10^-17^] and did not significantly differ in amplitude [*t*(3158) = 0.042, *p* = 0.966), Fig. 10C]. A six-cone model best fit adult control datasets (Vu358-UV, Vb415-B1, Vg460-G1, Vg480-G3, Vr556-R2, Vr575-R1). A four-cone model sufficed for *gnat2:trβ2* adults (Vb415-B1, Vg460-G1, Vr556-R2, Vr575-R1). Unlike *gnat2:trβ2* larvae (Fig. 9C), UV-cone signals were not significant in *gnat2:trβ2* adults (Fig. 10C).

At all levels of constant quantal stimulation long-wavelength spectral peaks between 560 and 570 nm are modeled for *gnat2:mYFP-2A-trβ2;mpv17^-/-^* (Fig. 10D). Models of the *mpv17^-/-^* control peak spectrally between 441 and 493 nm depending on stimulus brightness. Except at wavelengths greater than 540 nm, control amplitudes were greater than transgenic, due both to the greater cone sensitivities (lesser half-saturation irradiances), and to the significantly greater amplitudes of numerous mid- and short-wavelength cones (Vu358-UV, Vb415-B1, Vg460-G1, Vg480-G3).

As judged by residual variance no models were indistinguishable from the illustrated ones for *gnat2:mYFP-2A-trβ2;mpv17^-/-^* adults (*F*-tests for other models, *p* < 0.95). Model #235 was indistinguishable from model #219 (the best model) for the *mpv17^-/-^* control adults [*F*(0.9978, 1603, 1603) = 0.482, *p* = 0.96]. Models #235 and #219 both employed 6 cones, but model #235 substituted the G4 (Vg500) for the G3 (Vg480) cone.

### The larval dual opsin UV cone in trβ2 gain-of-function transgenics

Many cones in *gnat2:mYFP-2A-trβ2;mpv17^-/-^* larvae express mixed opsins, overlaying a red opsin on the native expression of a shorter-wavelength opsin (Fig. 7B). A few dual-opsin cones also occur in *crx:mYFP-2A-trβ2* larvae (Fig. 3B). The spectral physiology and molecular development of such cones is of interest. The wide spectral separation of red and UV opsins makes the two opsins in UV cones amenable to separate stimulation. One proposal is that zebrafish transgenic dual-opsin cones are analogous to the stable *mws-sws* ‘dual physiology’ cone configuration seen in rodents (Jacobs et al., 1991; Applebury et al., 2000; Nikonov et al., 2005; Suzuki et al., 2013). Another proposal is that the mixed-opsin cones in *gnat2:trβ2* zebrafish are transitional states, later to develop into red cones. The tendency to lose UV opsin signals in trβ2 gain-of-function transgenics suggests an adverse action on UV opsins, and a prominent model in mouse development is that trβ2 changes differentiated S cones to M cones (Swaroop et al., 2010).

To observe the effects of excess trβ2 on *sws1* (UV opsin) gene activity, UV opsin immunoreactivity was examined in double transgenic *gnat2:trβ2* larvae to which a fluorescent *sws1* opsin reporter gene was added (*gnat2:mYFP-2A-trβ2;sws1:nfsBmCherry;mpv17^-/-^*). Confocal micrographs of tangential planes through the cone photoreceptor layers in control and the double transgenic larvae show that at 5dpf (Fig. 11D, 11H) red-opsin immunoreactive cones are about twice as dense in *gnat2:trβ2* larvae (148,000 ±12,000 mm^-1^) as in controls [60,000 ±14,000 mm^-1^; *t*(6) = 5.52, *p* = 0.0015], but at this stage, the density of UV-opsin immunoreactive cones (Fig. 11B, 11F) is not affected *[gnat2:trβ2:* 48,000 ±3400 mm^-1^; control: 52,000 ±4100 mm^-1^; *t*(6) = 0.77, p = 0.47].

**Figure 11.**
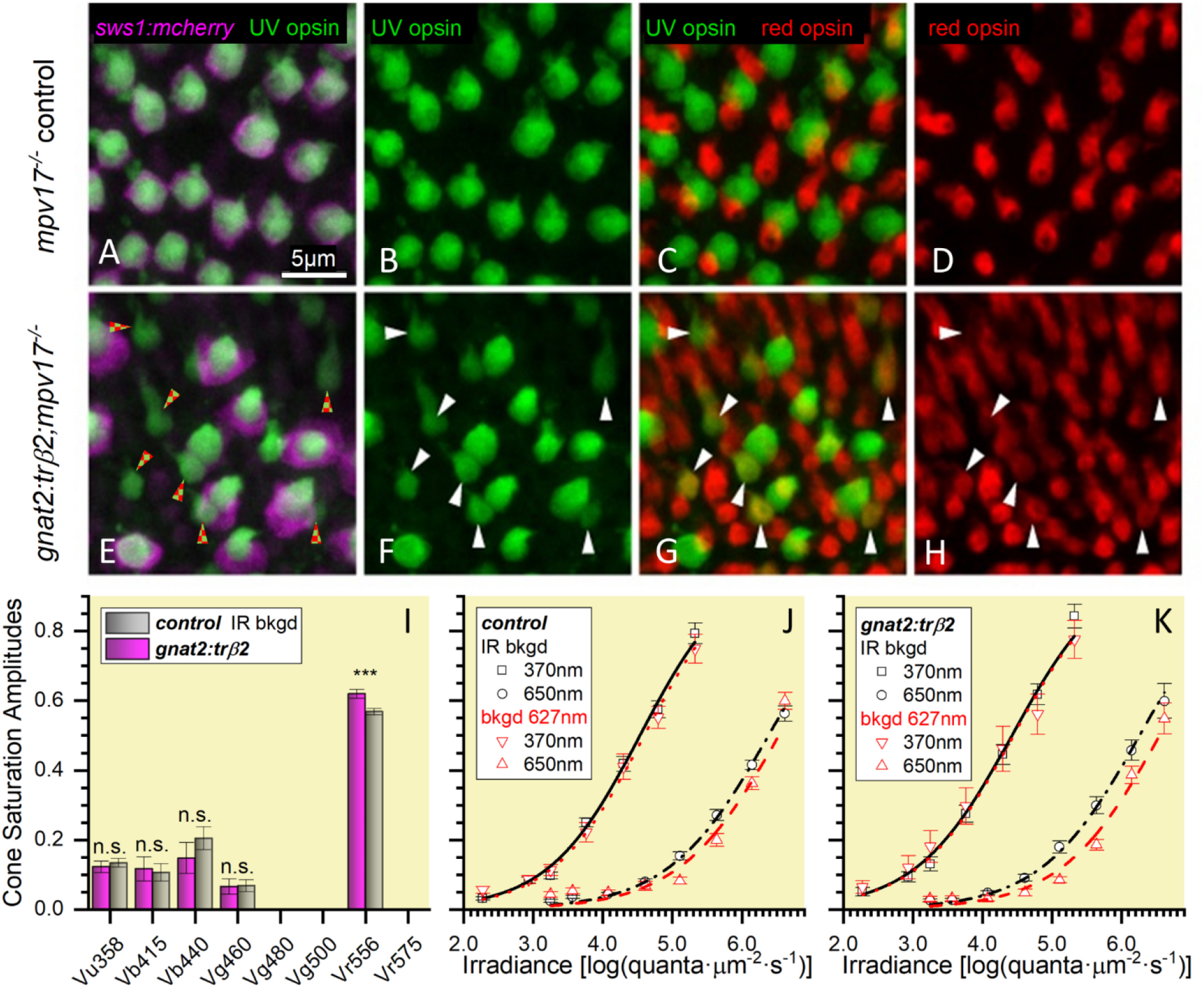
UV-opsin reporter inactivated in cones co-expressing UV and red opsins but red opsins produce no signal. ***A***, UV-opsin (SWS1) immunoreactivity (green) and florescence of a reporter transgene for *sws1* (*sws1:nfsBmCherry*, magenta) colocalize in *mpv17^-/-^* control UV cones. ***B***, UV-opsin immunoreactivity in control retinas. ***C***, In the control retina UV-opsin (green) and red-opsin immunoreactivity (red) localize in separate cones. ***D***, Red-opsin immunoreactivity in the control retina. ***E***, In *gnat2:trβ2* cones, UV-opsin immunoreactivity (green) is always found when there is *sws1* reporter gene fluorescence (*sws1:nfsBmCherry*, magenta), but not all UV-opsin immunoreactive cones show reporter-gene fluorescence (red and green checkered arrowheads). ***F***, UV-opsin immunoreactive cones in the *gnat2:trβ2* retina. ***G***, Co-expression of UV-opsin (green) and red-opsin (red) immunoreactivity in *gnat2:trβ2* cones. White arrowheads (***F***, ***G***, ***H***) point to cones expressing both UV (green) and red opsins (red). The dual opsin cones are the UV-opsin immunoreactive cones in ***E***, that lack *sws1* reporter fluorescence (red and green checkered arrowheads). ***H***, Red opsin immunoreactive cones in *gnat2:trβ2* retina. ***I***, Saturation amplitudes (*V_mi_* in Eq. 1, Fig. 2A) of signals from cone types in control (grey) and *gnat2:trβ2* (magenta) eyes. Control: Spectral algorithm fit to 1890 datapoints from 27 datasets recorded in 15 eyes; *gnat2:trβ2*, 1260 datapoints from 18 datasets recorded in 11 eyes. In each case the optimal fit was model #79. Asterisks give significance of differences (two sample *t*-tests, n.s., not significant) in cone-signal amplitudes [Vu358: *t*(3091) = 0.56, *p* = 0.57); Vb415: *t*(3091) = 0.25, *p* = 0.81); Vb440: *t*(3091) = 1.05, *p* = 0.29); Vg460: *t*(3091) = 0.084, *p* = 0.93); Vr556, *t*(3091) = 3.39, *p* = 0.0007)]. ***J***, Control strain (*mpv17^-/-^*) irradiance-response curves at 370 and 650 nm in the presence of infrared (IR) or 627-nm (red) backgrounds (bkgd). Control points are means ±SE (370 nm stimulus, IR background, *n*= 27; 370 nm stimulus, red background, *n* = 15; 650 nm stimulus, IR background, *n* = 54 or 27; 650 nm stimulus, red background, *n* = 30 or 15). ***K***, *gnat2:trβ2* irradiance-response curves at 370 and 650 nm in the presence of IR or red backgrounds. Points are means ±SE (370 nm stimulus, IR background, *n* = 18; 370 nm stimulus, red background, *n* = 12; 650 nm stimulus, IR background, *n* = 18 or 36; 650 nm stimulus, red background, n = 12 or 24). ***J***, ***K***, Hill function curves fit at each background and wavelength are constrained to have equal maximal amplitudes and coefficients. Cone-PIII ERG signals isolated with 20 mM Aspartate. ***A***-***K***, Eyes and retinas are from 5-dpf larvae.

For the *mpv17^-/-^* control, all UV-opsin immunoreactive cones (green) express the *sws1:nfsBmCherry* UV-opsin gene reporter (magenta) as an inner segment halo surrounding the narrower bright green immunofluorescence of the UV-cone outer segment (Fig. 11A). UV and red opsin immunoreactivities are segregated, being expressed in separate cone cells (Fig. 11C), the native larval opsin expression pattern (Allison et al., 2010). In the larval *gnat2:trβ2* double transgenic not all UV-opsin immunoreactive cones show the magenta halo of the *sws1:nfsBmCherry* reporter gene (Fig. 11E). In some, the *sws1* reporter gene generates no fluorescence, suggesting the native *sws1* gene locus is inactive, and UV-opsin, while still present, is no longer being synthesized. Only legacy SWS1 immunoreactivity remains. In figure 11G patterns of *gnat2:trβ2* immunoreactivity for UV-opsin and red-opsin are compared. The arrowheads point to cone cells where both UV and red opsin are co-expressed. A comparison of figure 11F with figure 11E reveals that UV-opsin immunoreactive cones with inactive *sws1* reporter genes are the same cones that are double immunoreactive for both UV- and red-opsins (Fig. 11E, checkered arrowheads). The expression of red opsin in a UV cone by the gain-of-function *gnat2:trβ2* transgene appears incompatible with continued expression of UV-opsin, suggesting that the overabundance trβ2 in a UV-cone either directly or indirectly blocks the *sws1* gene. Therefore, co-expression of UV-and red-opsin immunoreactivity cannot continue indefinitely, as in rodents, but would be limited by the catabolism of previously expressed, but not renewed, UV-opsin.

At the same embryonic developmental stage (5 dpf), the functional properties of UV-cones with co-expression of red opsin were explored. In this earliest embryonic dataset, the same 5-cone model (#79) was the best fit for both control and *gnat2:trβ2* gain-of-function larvae (Fig. 11-I), and the residual variance of the *gnat2:trβ2* dataset was not significantly greater when modeled with the control amplitude distribution [*F*(1.031,1230,1235) = 0.705, *p* = 0.591], although the R2-cone saturation amplitude (Vr556) trended greater (9%) in *gnat2:trβ2* [*t*(3091) = 3.39, *p* = 0.0007)]. Saturation amplitudes of shorter-wavelength cone signals (Vu358-UV, Vb415-V1, Vb440-B2, Vg460-G1) in *gnat2:trβ2* and control were not significantly different. UV-cone saturation amplitude at 5 dpf appeared unaffected by numerous of its members both lacking *sws1* gene activity and co-expressing red opsin.

Red chromatic adaptation should desensitize UV/red mixed opsin cones to UV stimulation. 650-nm red stimuli are not absorbed by UV opsins, but 370-nm UV stimuli are strongly absorbed. We generated irradiance-response functions at 650 nm and 370 nm in the presence of red or IR backgrounds (Figs. 11J, 11K). For 370-nm stimuli, the control, irradiance-response curves overlap in both datapoints and Hill-function curve fits regardless of red or IR background illumination (Fig. 11J). Hill function fits give semisaturation values of 4.53 log(quanta·μm-^2^·s^-1^) for the IR background and 4.59 for the 627-nm background, not distinguishable [*t*(290) = 1.26, *p* = 0.210)], an expected result as UV-opsin does not absorb either the 627-nm or the IR adapting light. As the 650-nm stimuli (Fig. 11J) are not as efficiently transduced, even by red cones, the 650-nm irradiance-response functions shift to greater irradiances. On the IR background the 650-nm semi-saturation irradiance is 6.38 log(quanta·μm-^2^·s^-1^) but increases to 6.53 log(quanta·μm- ^2^·s^-1^) with the 627-nm background, a significant desensitization [*t*(583) = 3.66, *p* = 0.0003)]. The absorption of the red background by the red-cone red opsin reduces red-cone sensitivity (Fig. 11J).

What was unexpected was that the 370-nm irradiance functions for the *gnat2:trβ2* gain-of-function larvae, with co-expression of red opsin in some UV cones, were unaffected by the red background (Fig. 11K). The red opsin in the mixed-opsin UV cones, which would be activated by the 627-nm background, does not desensitize 370-nm UV signals. For 370-nm stimuli, Hill fits give semi-saturation irradiances of 4.42 log(quanta·μm-^2^s^-1^) on the IR background and 4.44 log(quanta·μm-^2^s^-1^) on the 627-nm background, values not significantly different [*t*(205) = 0.300, *p* = 0.765)]. But the same red background significantly desensitizes red-opsin signals from *gnat2:trβ2* red cones, as seen with 650-nm stimuli (Fig. 11K). The Hill semi-saturation changed from 6.26 log(quanta·μm^-2^·s^-1^) on the IR background to 6.55 log(quanta·μm^2^·s^-1^) on the 627-nm background, a significant sensitivity loss [*t*(415) = 5.01, *p* = 1.5× 10^-12^], which demonstrates the effectiveness of this background for red opsins (Fig. 11K). The failure of long-wavelength backgrounds to affect UV signals from mixed-opsin cones has also been observed in ERG spectra of rodents (Jacobs et al., 1991). In zebrafish the spectral physiology of UV cones appears, at least early in development at 5 dpf, not to be affected by the introduction of transgenic red opsins into many UV-cone members.

### Thyroxin receptor β2 gain-of-function transgenes alter cone morphology

Thyroid hormone receptor β2 is required in zebrafish for both the development of red cones and the expression of red opsins (Deveau et al., 2020). Morphologically, adult red cones are the principal members of zebrafish double cones (Engstrom, 1960; Raymond et al., 1993). Double cones failed to develop in *trβ2^-/-^* mutants (Deveau et al., 2020). The impact of trβ2 in determining the course of cone morphological development is seen in transverse optical sections of individual transgenic cones from *in vivo* 6-dpf larvae (Fig. 12). These are higher magnifications taken from larval retina confocal image stacks such as seen in figures 1D, 1E and 1F. On inspection, the fluorescent-reporter shapes of trβ2 gain-of-function cones are distinct from the control morphologies of red, blue and UV cones, the latter marked by the reporter fluorescent proteins expressed in *sws2:GFP* (blue cones) and *sws1:GFP;trβ2:tdTomato*(UV and red cones) (Fig. 1D, 12A, 12C).

**Figure 12.**
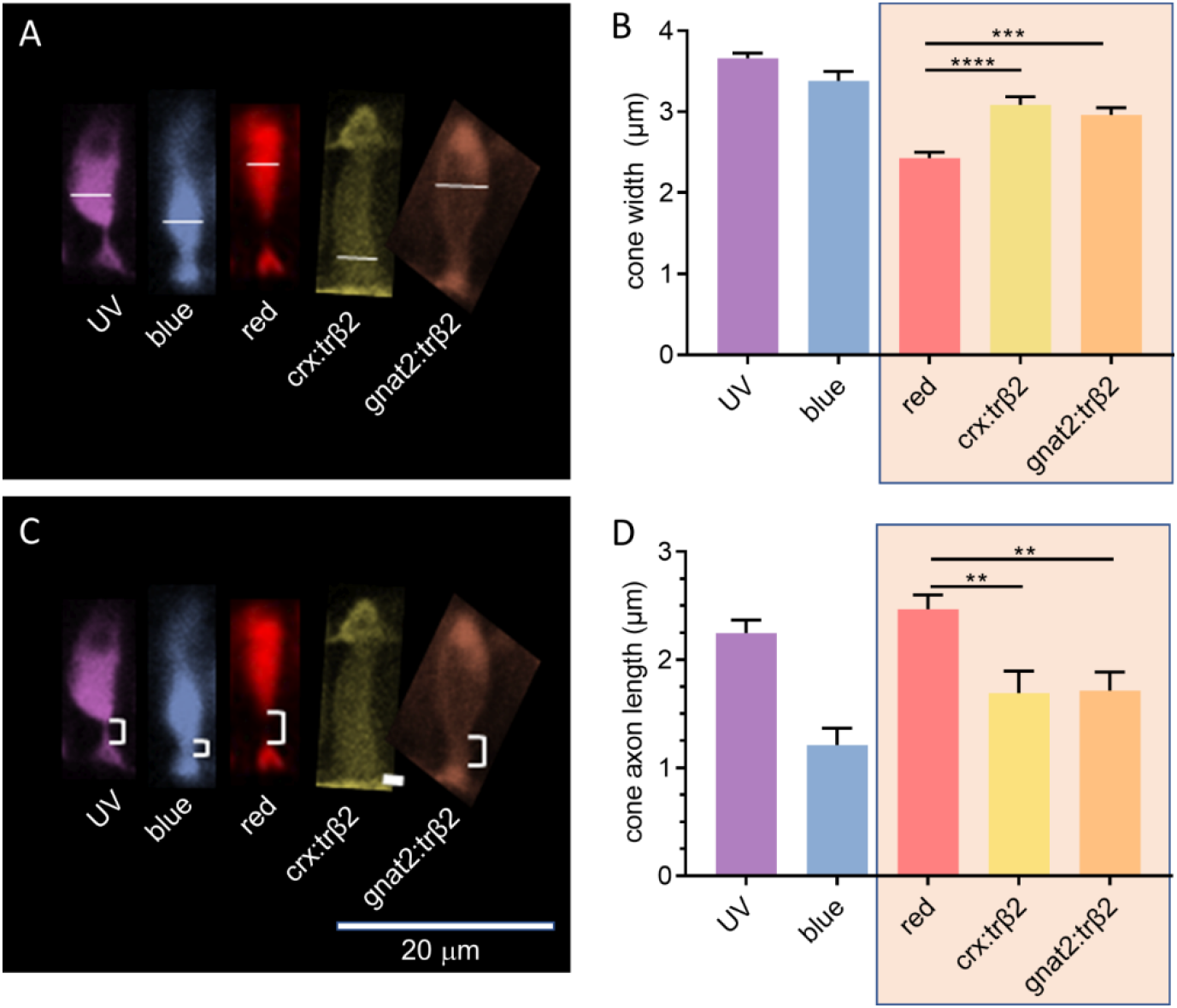
Thyroxin-receptor-β2 gain-of-function transgenes alter cone morphology. ***A***, Width of cone types identified by transgene markers, is measured at the greatest extent of the inner segment. ***B***, The *trβ2* gain-of function cones are significantly wider than red-cones. ***C***, The cone axon length is measured from the base of the inner segment to the apex of the cone pedicle. ***D***, The *trβ2* gain- of-function axon lengths are significantly shorter than red-cone axon lengths. ***C***, ***D***, Asterisks indicate significant differences (Graphpad convention, ANOVA and tukey-post-hoc *p*-values given in text). ***A***,***C***, Images are of 6-dpf *in-vivo* larval fluorescent cones from confocal stacks. Control UV and red cones were imaged in *sws1:GFP;trβ2:tdTomato* larvae; blue cones, in *sws2:GFP* larvae; fluorescent *trβ2* gain-of-function cones, in *crx:mYFP-2A-trβ2* and *gnat2:mYFP-2A-trβ2;mpv17^-/-^* larvae. Larvae were anesthetized with MS222 and embedded in agarose after raising to 6 dpf in 300 μM PTU to block melanin formation in the pigment epithelium.

The maximal widths of the five sorts of cone inner segments illustrated differed [ANOVA, *F*(4, 356) = 36.98, *p* < 0.00001]. The maximal widths for cone inner segments of *crx:mYFP-2A-trβ2* labeled cones are significantly wider than red cones (*post-hoc Tukey, p* < 0.000001), as are the maximal widths of cone inner segments of *gnat2:mYFP-2A-trβ2;mpv17^-/-^* labeled cones (*post-hoc Tukey, p =*.00024). Neither trβ2 gain-of-function cone is as wide as the inner segments of UV cones (*post-hoc Tukey: crx:mYFP-2A-trβ2, p*= 0.00005; *gnat2:mYFP-2A-trβ2;mpv17^-/-^,p* < 0.000001). The inner segment widths of the *crx:mYFP-2A-trβ2* cones were not distinguishable from blue cones (*post-hoc Tukey: p* = 0.126), while *gnat2:mYFP-2A-trβ2* widths were marginally narrower than blue cones (*post-hoc Tukey:p* = 0.00764).

The lengths of axons connecting the base of cone inner segments and the apex of cone synaptic pedicles differed among the cone types illustrated in figure 12C [ANOVA, *F*(4, 290) = 9.692,*p* < 0.00001]. In the trβ2 gain-of-function cones, the length of the axon is significantly shorter than in red cones (*post-hoc Tukey: crx:mYFP-2A-trβ2, p* = 0.00644; *gnat2:mYFP-2A-trβ2;mpv17^-/-^*,*p* = 0. 00883, Fig. 12D). The lengths are not distinguishable from blue cone axons (*post-hoc Tukey: crx:mYFP-2A-trβ2,p* = 0.21190; *gnat2:mYFP-2A-trβ2;mpv17^-/-^*,*p* = 0. 17500, Fig. 12D) or UV cones axons (*post-hoc Tukey: crx:mYFP-2A-trβ2, p* = 0. 10775; *gnat2:mYFP-2A-trβ2;mpv17^-/-^, p* = 0. 13379, Fig. 12D).

On morphometrics, the mYFP-marked trβ2 gain-of-function cones have not become red cones at 6 dpf, but nonetheless enhance red cone signal amplitudes. Gain-of-function trβ2 morphologically alters cones and appears to shift the metrics towards a larval blue-cone pattern.

### Impact of excess thyroid hormone receptor β2 on development of zebrafish spectral signals

During zebrafish maturation, patterns of cone opsin mRNA expression change through interactions with a ‘developmental factor’ (Takechi and Kawamura, 2005). Green- and red-cone spectral peaks shift from shorter to longer wavelengths by adulthood (Nelson et al., 2019) as different members of gene-duplicated green (*Rh2*) and red (*lws*) opsin groups are sequentially expressed. In the present control data, the R2 opsin signal (Vu556) is largest in embryonic (5 dpf, 6dpf) and juvenile (12 dpf) ages but the R1 opsin signal (Vr575) attains equal amplitude status in adults, while, over the same developmental course, the UV cone signal (Vu358) diminishes (Fig. 13A, 13B, WT and *mpv17^-/-^* controls, gray bars). Introduction of gain-of-function trβ2 shifts native developmental patterns of opsin expression. In the adults of both *crx:trβ2* and *gnat2:trβ2* gain-of-function transgenics, UV- (Vu358), blue- (Vb415, Vb440) and green- (Vg460, Vg480, Vg500) cone signals are reduced or extinguished by adulthood. In embryos and juveniles mainly R2 signals (Vr556) increase but in adults, R1 signals (Vr575) increase. (Fig. 13A, 13B). In *crx:trβ2* transgenics, as early as 5 dpf UV- (Vu358) and B1- (Vb415) cone signals decrease significantly, to 11% and 20% of WT control, while R2-cone amplitude (Vr556) increases 230% (Fig. 13A). In *crx:trβ2*, the altered pattern of signal amplitudes from cone types persists throughout development, with UV cone signals becoming undetectable from 6 dpf on through adulthood (Fig. 13A, 6dpf, 12 dpf, Adult) and there is a failure to detect blue and green cone signals after the 12-dpf juvenile stage (Fig. 13A, Adult). Signals from red cones increase throughout embryonic (5 dpf, 6 dpf), juvenile (12 dpf) and adult developmental stages. R2 cone amplitudes (Vr556) increase at both 5 dpf and 12 dpf [12 dpf: *t*(1558) = 6.29, *p* = 3.9 × 10^-10^] and R1 cone signals (Vr575) increase for 6-dpf embryos and for adults [6 dpf: *t*(1825) = 2.72,*p* = 0.0067; Adult: *t*(2818) = 3.92,*p* = 9.2 × 10^-5^; Fig. 13A].

**Figure 13.**
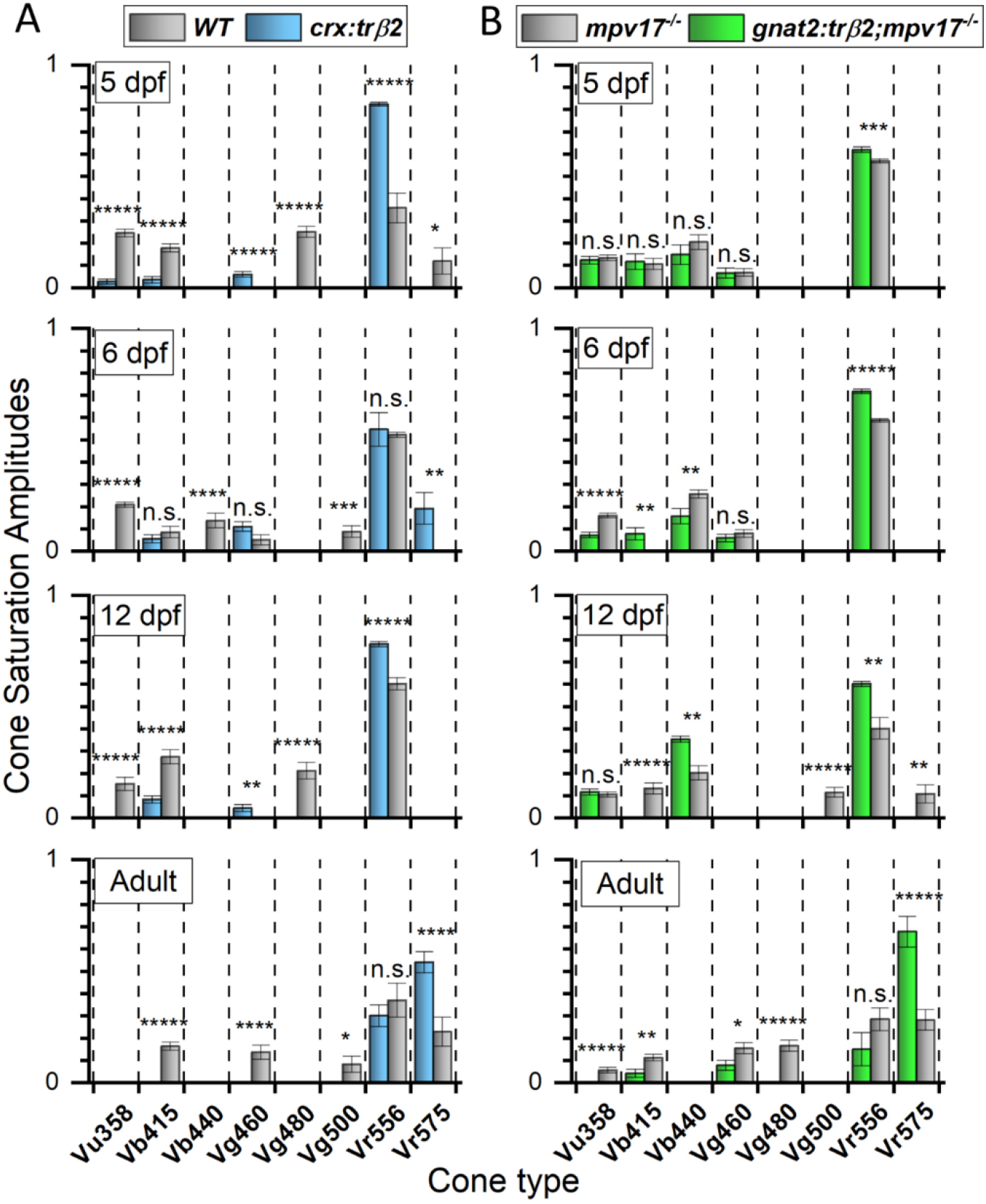
Signal development in red-, green-, blue- and UV-cones. ***A***, Throughout development red-cone signals (Vr575, Vr556) are larger than WT in *crx:mYFP-2A-trβ2* while blue-, green- and UV-cone signals (Vb415, Vb440; Vg460, Vg480, Vg500; Vu360) are smaller, disappearing with time. ***B***, The overall signal amplitudes of red, green and blue cones in *gnat2:mYFP-2A-trβ2;mpv17^-/-^*are indistinguishable from *mpv17^-/-^ (roy orbison*) controls at 5 dpf but red-cone amplitudes increase while green-, blue- and UV-cone signals decrease later in development. ***A***, ***B***, ‘5 dpf’ and ‘6 dpf are larval or embryonic stages. ‘12 dpf is a juvenile stage and ‘Adult’ is 8-18 months. In each bar chart spectral datasets from multiple eyes are combined and fit by the equation 1 (Fig. 1) algorithm to model the contributing cone types and their saturation voltages. A cone only detected in one of the two strains is evaluated by a one-sample *t*-test and a cone detected in both strains by a two-sample *t*-test. Asterisks denoting significance (Graphpad convention, n.s., not significant) either compare detected transgenic and control amplitudes for a cone type or show the significance of the single detected amplitude. ***A***, <u>5 dpf</u>, 7 WT eyes, 11 *crx:trβ2* eyes: *Vu358* (UV), *t*(2465) = 10.4*, p* = 5.8 × 10^-25^; *Vb415* (B1), *t*(2465) = 5.74, *p* = 1.1 × 10^-8^; *Vg460* (G1), *t*(1638) = 4.54, *p* = 6.0 × 10^-6^; *Vg480* (G3), *t*(827) = 10.3, *p* = 1.6 × 10^-23^; *Vr556* (R2), *t*(2465) = 9.51, *p* = 4.3 × 10^-21^; *Vr575*, (R1), *t*(827) = 2.02, *p* = 0.043. 6 dpf. 10 WT eyes, 11 *crx:trβ2* eyes: *t*-tests given in figure 5. 12 dpf. 3 WT eyes, 6 *crx:trβ2* eyes. *Vu358* (UV), *t*(243) = 5.12, *p* = 6.2 × 10^-7^; *Vb415* (B1), *t*(1558) = 4.86, *p* = 1.3 × 10^-6^; *Vg460* (G1), *t*(1315) = 2.86, *p* = 0.00435; *Vg480* (G3), *t*(243) = 5.61, *p* = 5.5 × 10^-8^; *Vr556* (R2), *t*(1558) = 6.30, 3.90 × 10^-10^. Adult. 14 WT eyes, 16 *crx:trβ2* eyes: *t*-tests given in figure 6. ***B***, 5 dpf. 15 *mpv17^-/-^* eyes, 11 *gnat2:trβ2;mpv17^-/-^* eyes: *t*-tests given in figure 11.6 dpf. 15 *mpv17^-/-^* eyes; 17 *gnat2:trβ2; mpv17^-/-^* eyes: *t*-tests given in figure 9. 12 dpf. 16 *mpv17^-/-^* eyes; 9 *gnat2:trβ2;mpv17^-/-^*eyes: *Vu358* (UV), t(2540) = 0.59, *p* = 0.55; *Vb415* (B1), *t*(1786) = 5.31, *p* = 1.2 × 10^-7^; *Vg440* (B2), *t*(2540) = 3.04, *p* = 0.0023; *Vg500* (G4), *t*(1786) = 5.22, *p* = 2.0 × 10^-7^; *Vr556* (R2), *t*(2540) = 2.65, *p* = 0.00815; *Vr575* (R1), *t*(1786) = 2.66, *p* = 0.0079. Adult. 13 *mpv17^-/-^* eyes; 11 *gnat2:trβ2;mpv17^-/-^* eyes: *t*-tests given in figure 10.

For the *gnat2* promoter, the consequences of gain-of-function trβ2 are more gradual in developmental course (Fig. 13B). In the 5dpf *gnat2:mYFP-2A-trβ2* embryonic larvae, the overall distribution of signal amplitudes from cone types is not significantly different from those found in the control (*mpv17^-/-^*)siblings, but R2 cone signals (Vr556) trend larger (Fig. 13B, 5dpf). At 6dpf, cone signal distribution differs significantly and enhancement of R2 amplitudes increases [6 dpf: 22%, *t*(3422) = 9.42, *p* = 5.6 × 10^-23^]. In 12 dpf juveniles, R2 amplitude increases by 50% [*t*(2540) = 2.65, *p* = 0.0082], but in adults red-cone enhancement switches from R2 (Vr556) to R1 (Vr575), where a 241% enhancement is seen [*t*(2953) = 4.86, *p* = 1.2 × 10^-6^], with no significant change seen in R2-cone amplitudes [*t*(2953) = 1.52, *p* = 0.127]. In *gnat2:trβ2*, UV-cone signals (Vu358) persist through larval and juvenile stages, with amplitudes either not significantly different [5 dpf: *t*(3091) = 0.563, *p* = 0.574; 12 dpf: *t*(2540) = 0.594, *p* = 0.553] or less than control [6 dpf: 45%, *t*(3422) = 5.36, *p* = 8.7 × 10^-8^], but unlike the significant UV signals of control adult [*t*(1579) = 4.81,*p* = 1.7 × 10^-6^], no UV-signal was detected in *gnat2:trβ2* adults (Fig. 13B). B1- and G1- cone signals persist in *gnat2:trβ2* adults but are significantly reduced in amplitude as compared to controls [B1: 37%, *t*(2953) = 2.97, *p* = 0.0030; G1, 50%, *t*(2953) = 2.23, *p* = 0.024].

## Discussion

Modeling of the massed cone signals from zebrafish retinas yielded estimates of amplitude contributions from eight spectrally distinct cone types during embryonic, juvenile, and adult developmental stages. The technique provided a window on the impact of transgene-induced overabundance in the red-cone transcription factor trβ2 for the balance among electrical signals from red-cone opsins and all other opsin types. Two trβ2 gain-of-function transgenics were studied. In *crx:mYFP-2A-trβ2*, the *crx* promoter introduced trβ2 into retinal progenitor cells, whether ultimately fated to become cone cells or other types, such as bipolar cells (Shen and Raymond, 2004). In *gnat2:mYFP-2A-trβ2;mpv17^-/-^*, the *gnat2* promoter increased trβ2 levels only in differentiated cone types, including green, blue and UV cones where it is not native, as well as adding an extra dose to the red-opsin cones where it is normally expressed (Ng et al., 2001; Suzuki et al., 2013). Neither transgene caused major alterations in the amplitudes or kinetics of massed cone signals as isolated from the ERG by blockade of cone synapses. In *crx:mYFP-2A-trβ2* larvae, response amplitudes were larger than WT controls, as was the variance in amplitudes. In the adults, peak times of both onset and offset waveform elements were significantly faster. Neither alteration suggested a major net influence on phototransduction. In *gnat2:mYFP-2A-trβ2;mpv17^-/-^* there were no significant changes either in amplitudes or in onset and offset kinetics. But in both transgenics, significant changes were found in the relative contributions from different cone types to retinal spectral responses.

### Opsin signals in *crx:trβ2* transgenics

For *crx:trβ2* transgenics, the net amplitude of red opsin signals increased at all developmental stages, as did the densities of red-opsin immunoreactive cones, as seen previously by Suzuki et al (2013), suggesting increased cone numbers led to increased signal strength. But LWS2 (R2) opsin signals were favored in larvae and juveniles, while LWS1 (R1) signal amplitudes became disproportionately large in adults. The latter suggests that, in addition to augmentation of red cone abundance and signal amplitude, excess trβ2 favored R1 opsin expression but to do so required a cofactor expressed only late in development. Individual zebrafish red cones express either one or the other of the *lws1* or *lws2* opsin genes but not both (Tsujimura et al., 2010) and thyroid hormone, the trβ2 ligand, can induce a cone to switch opsins from LWS2 to LWS1 (Mackin et al., 2019). In situ hybridization indicates that red cones in embryonic and larval zebrafish choose mainly the *lws2* opsin gene, while adult red cones are more likely to transcribe *lws1*, particularly in peripheral retina (Takechi and Kawamura, 2005). The WT and *mpv17^-/-^*control cone-ERG spectral analyses support a direct proportionality to the red-opsin transcript and expression developmental profiles. LWS2 (R2) opsin signals were the largest, and most often the only, red-cone signals detected in embryos and juveniles of control animals, but about equal amplitudes of both R1 and R2 signals appeared in adult controls. In gain-of-function *crx:trβ2* larvae and juveniles, with one exception, it was similarly R2 signals that trβ2 overabundance increased. In *crx:trβ2* adults, it was R1 signals that increased, becoming the largest-amplitude cone spectral signal, and shifting cone ERG spectral peaks from mid-spectrum in controls to long wavelengths in *crx:trβ2*. Augmentation of LWS1 transcript occurs in adult *six7* transcription factor knockouts (Ogawa et al., 2015), suggesting a model where this transcription factor inhibits *lws1*, but might itself be inhibited by trβ2. Alternatively thyroid hormone has been shown to favor expression of LWS1 over LWS2 transcript (Mackin et al., 2019) and a different model would propose greater thyroid hormone levels in adults would lead to greater levels of bound trβ2, proportionally favoring a switch to LWS1 cone physiology.

Zebrafish UV (SWS1) cones are the molecular phylogenetic relatives of mammalian S-cones (Terakita, 2005). In *crx:trβ2* transgenics the amplitude of UV cone signals was greatly decreased or eliminated at all developmental stages, suggesting the diminished densities of UV opsin immunoreactive cones noted herein and by Suzuki et al (2013), led directly to decreased UV-cone signal strength. With the progression of developmental stages in teleosts, the role of UV cones typically diminishes (Cheng et al., 2006; Carleton, 2009; Nelson et al., 2019). Juvenile trout lose UV sensitivity as they mature. The process is thyroid hormone sensitive (Browman and Hawryshyn, 1994) and correlates with regional loss of SWS1 UV-opsin immunoreactivity and UV cone morphologies. Based on regional maturation in thyroid hormone and thyroid hormone receptor levels within retinal quadrants, together with experimental treatments with thyroid hormone (T4), Raine and Hawryshyn (2009) proposed that regional loss of UV-cone signaling was caused by regional increases in both in T4 and trβ. In zebrafish, similarly, T4 reduced UV-opsin transcript levels (Mackin et al., 2019). Present results add that increased levels of trβ2 receptor itself reduce the signal amplitude and numbers of UV cones and that trβ2 is a potential candidate regulating their density. How this might occur without a transgene is less clear, as trβ2 is not normally expressed by UV cones (Suzuki et al., 2013).

### Opsin signals in *gnat2:trβ2* transgenics

For *gnat2:trβ2* 5-day embryos, introduction of trβ2 into functional cones of all spectral types doubled the numbers of red-opsin immunoreactive cones without changing the densities of cones with other opsin immunoreactivity, a result first noted by Suzuki et al (2013) and repeated here. But in the present cone ERG analysis, red-cone signal amplitude increased less than 10 percent. In this counterexample the densities of red opsin immunoreactive cones and signal strength were not proportional. Densities of green-, blue- and UV-opsin immunoreactive cones, and the distribution of opsin signals among them, were unchanged. As discovered by Suzuki et al (2013) and confirmed here, much of the increase in density for red opsin immunoreactive cones is accounted for by co-expression. Suzuki et al (2013) found red-green and red-UV immunoreactive cones. To this we add cones immunoreactive for both red and blue opsins. Red opsins induced in differentiated *gnat2:trβ2* green, blue and UV cones evidently do not immediately result in a greater red-cone electrical signal at the 5-day larval stage. By adulthood the spectral signals *gnat2:trβ2* do come to resemble those of adult *crx:trβ2* red opsin dichromats, with largest amplitudes originating from LWS1 opsins and green blue and UV amplitudes reduced or suppressed. This suggests the introduction of trβ2 into differentiated cones quickly expresses red opsins but is much slower to generate electrical signals than is the case with *crx:trβ2*, which introduces trβ2 into cone progenitors.

In embryonic *gnat2:trβ2* transgenics the amplitude of UV cone signals was not affected by co-expression of red opsins. To examine red-UV immunoreactive cones further we examined *sws1* reporter gene expression and the sensitivity of UV cones to adaptation by red backgrounds. In controls, *sws1* reporter activity and UV-opsin immunoreactivity co-localized, but in *gnat2:trβ2* transgenics UV-opsin immunoreactive cones that co-expressed red opsin lost *sws1* reporter activity, suggesting that, even without transcriptional resupply of UV opsin, long-lived previously synthesized UV-opsin survived in the cone disks and functioned for some time after synthesis was suppressed. We suggest that the co-expressed red-opsins remain electrically dormant during this initial period, as red backgrounds which desensitize red-cone signals failed to desensitize UV-cone signals in 5-dpf *gnat2:trβ2*. In the longer term UV-opsin would be lost to disk shedding (O’day and Young, 1978). This may be a path by which UV/red mixed opsin cones are gradually lost, and/or converted to red cones, similar to an important model in mouse M-cone embryogenesis from primordial UV cones (Swaroop et al., 2010).

### Suppression of green and blue cones

Overproduction of trβ2 in zebrafish gain-of-function transgenics reduced or eliminated green-cone (Rh-2) and blue-cone (SWS2) signals. Vb415 (B1, SWS2) and Vg460 (G1, Rh2-1) signals were significant in *crx:trβ2* embryos and juveniles but lost in adults. G1-cone signals might be either increased or decreased compared to controls in *gnat2:trβ2* embryos or juveniles but both were reduced in adults. In adult zebrafish *trβ2^-/-^* mutants (Deveau et al., 2020) green-cone signals increased significantly in amplitude. These observations suggest a late-stage inhibitory effect of trβ2 on green and blue cones. Knockout of the zebrafish homeobox transcription factor *six7* eliminates green-cone Rh2 transcripts in adults and *six6* knockouts adversely affected blue-cone SWS2 transcript (Ogawa et al., 2015; Ogawa et al., 2019). An association of these transcription factors with blue and green cones was made by single-cell sequencing and machine learning methods (Ogawa and Corbo, 2021). Speculatively there is an inhibitory action of trβ2 on these homeobox genes.

### Changes in cone morphology

Larval cone morphologies in both trβ2 gain-of-function transgenics were altered. In *gnat2:mYFP-2A-trβ2*,where the transgene was expressed in all cone types, larval cones expressing the mYFP fluorescent transgene reporter did not look like red cones, resembled none of the control cone-type morphologies, but had a transgenic shape closest in morphometrics to, but visually distinguishable from, the control larval blue cones. There appears to be an early alteration of native morphologies induced by activity of the gain-of-function transgene. In *crx:mYFP-2A-trβ2* the mYFP transgene reporter was more sparsely expressed in the cone layer, nonetheless revealing altered cone morphology with inner segment widths and axon lengths similar to *gnat2:mYFP-2A-trβ2* cones. The basis of these trβ2-induced shape changes is not known, but despite them, the cones are robustly functional. Clearly trβ2 is importantly involved in large swaths of cone development, including the formation of adult double cones, with characteristic Arrestin 3a antigenicity (Deveau et al., 2020). Thyroid hormone receptor β2 pathways are not yet fully elaborated, but studies of zebrafish mutants and gain-of function transgenics expand the inventory of physiological, morphological, and genetic targets and provide insight into further roles in development.

## Acknowledgements

Current affiliations: **AB**, Department of Anatomy and Biology, George Washington University, Washington DC, 20052; **TS**, Washington University School of Medicine, St. Louis, MO, 63110; **LJE**, Neuroscience, Johns Hopkins Medical School, Baltimore, MD 21205; **TY**, Ophthalmology & Visual Sciences, Washington University School of Medicine, St. Louis, MO 63110; **SSP**, Center for Visual Science, University of Rochester, New York, 14627.

## Conflict of interest statement

Authors report no conflicts of interest

## Funding sources

This work was supported by The Intramural Program of the National Institutes of Neurological Disorders and Stroke, National Institutes of Health, Bethesda, Maryland and by National Institutes of Health Grant EY14356 (to Rachel O. L. Wong). TY research contributions conducted while in the ROL lab at the University of Washington.

## Notes

### Competing Interest Statement

The authors have declared no competing interest.

### Summary of Updates

Revisions of title, abstract, introduction, discussion, author affiliations and minor text edits.

## References

Allison WT, Barthel LK, Skebo KM, Takechi M, Kawamura S, Raymond PA (2010) Ontogeny of cone photoreceptor mosaics in zebrafish. Journal of Comparative Neurology 518:4182–4195.

Applebury M, Antoch M, Baxter L, Chun L, Falk J, Farhangfar F, Kage K, Krzystolik M, Lyass L, Robbins J (2000) The murine cone photoreceptor: a single cone type expresses both S and M opsins with retinal spatial patterning. Neuron 27:513–523.

Baylor D, Fuortes M (1970) Electrical responses of single cones in the retina of the turtle. The Journal of Physiology 207:77–92.

Bian W-P, Pu S-Y, Xie S-L, Wang C, Deng S, Strauss PR, Pei D-S (2021) Loss of mpv17 affected early embryonic development via mitochondria dysfunction in zebrafish. Cell death discovery 7:1–10.

Brockerhoff SE, Rieke F, Matthews Hr, Taylor MR, Kennedy B, Ankoudinova I, Niemi GA, Tucker CL, Xiao M, Cilluffo MC, Fain GL, Hurley JB (2003) Light stimulates a transducin-independent increase of cytoplasmic Ca2+ and suppression of current in cones from the zebrafish mutant nof. J Neurosci 23:470–480.

Browman HI, Hawryshyn CW (1994) The developmental trajectory of ultraviolet photosensitivity in rainbow trout is altered by thyroxine. Vision research 34:1397–1406.

Carleton K (2009) Cichlid fish visual systems: mechanisms of spectral tuning. Integrative zoology 4:75–86.

Cheng CL, Flamarique IN, Hárosi FI, Rickers-Haunerland J, Haunerland NH (2006) Photoreceptor layer of salmonid fishes: transformation and loss of single cones in juvenile fish. Journal of Comparative Neurology 495:213–235.

Chinen A, Hamaoka T, Yamada Y, Kawamura S (2003) Gene duplication and spectral diversification of cone visual pigments of zebrafish. Genetics 163:663–675.

Connaughton VP, Nelson R (2000) Axonal stratification patterns and glutamate-gated conductance mechanisms in zebrafish retinal bipolar cells. J Physiol 524:135–146.

Connaughton VP, Nelson R (2021) Ganglion cells in larval zebrafish retina integrate inputs from multiple cone types. Journal of Neurophysiology 126:1440–1454.

D’Agati G, Beltre R, Sessa A, Burger A, Zhou Y, Mosimann C, White RM (2017) A defect in the mitochondrial protein Mpv17 underlies the transparent casper zebrafish. Developmental Biology 430:11–17.

Dartnall H, JA (1953) The interpretation of spectral sensitivity curves. British Medical Bulletin 9:24–30.

Deveau C, Jiao X, Suzuki SC, Krishnakumar A, Yoshimatsu T, Hejtmancik JF, Nelson RF (2020) Thyroid hormone receptor beta mutations alter photoreceptor development and function in Danio rerio (zebrafish). PLoS genetics 16:e1008869.

Engstrom K (1960) Cone types and cone arrangement in the retina of some cyprinids.. Acta Zoologica 41:277–295.

Grant GB, Dowling JE (1995) A glutamate-activated chloride current in cone-driven ON bipolar cells of the white perch retina. J Neurosci 15:3852–3862.

Grant GB, Dowling JE (1996) On bipolar cell responses in the teleost retina are generated by two distinct mechanisms. J Neurophysiol 76:3842–3849.

Hughes A, Saszik S, Bilotta J, Demarco PJ, Jr., Patterson WF, 2nd (1998) Cone contributions to the photopic spectral sensitivity of the zebrafish ERG. Visual Neuroscience 15:1029–1037.

Jacobs GH, Neitz J, Deegan II JF (1991) Retinal receptors in rodents maximally sensitive to ultraviolet light. Nature 353:655.

Kennedy BN, Alvarez Y, Brockerhoff SE, Stearns GW, Sapetto-Rebow B, Taylor MR, Hurley JB (2007) Identification of a Zebrafish Cone Photoreceptor-Specific Promoter and Genetic Rescue of Achromatopsia in the nof Mutant. Invest Ophthalmol Vis Sci 48:522–529.

Mackin RD, Frey RA, Gutierrez C, Farre AA, Kawamura S, Mitchell DM, Stenkamp DL (2019) Endocrine regulation of multichromatic color vision. Proceedings of the National Academy of Sciences 116:16882–16891.

Marks W (1965) Visual pigments of single goldfish cones. The Journal of Physiology 178:14.

Nelson RF, Singla N (2009) A spectral model for signal elements isolated from zebrafish photopic electroretinogram. Visual Neuroscience 26:349–363.

Nelson RF, Balraj A, Suresh T, Torvund MM, Patterson SS (2019) Strain variations in opsin peaks in situ during zebrafish development. Visual neuroscience 36:E010.

Ng L, Hurley JB, Dierks B, Srinivas M, Saltó C, Vennström B, Reh TA, Forrest D (2001) A thyroid hormone receptor that is required for the development of green cone photoreceptors. Nature genetics 27:94–98.

Nikonov SS, Daniele LL, Zhu X, Craft CM, Swaroop A, Pugh Jr EN (2005) Photoreceptors of Nrl-/-mice coexpress functional S-and M-cone opsins having distinct inactivation mechanisms. The Journal of general physiology 125:287–304.

O’day WT, Young RW (1978) Rhythmic daily shedding of outer-segment membranes by visual cells in the goldfish. The Journal of cell biology 76:593–604.

Ogawa Y, Corbo JC (2021) Partitioning of gene expression among zebrafish photoreceptor subtypes. Scientific reports 11:1–13.

Ogawa Y, Shiraki T, Kojima D, Fukada Y (2015) Homeobox transcription factor Six7 governs expression of green opsin genes in zebrafish. Proc R Soc B 282:20150659.

Ogawa Y, Shiraki T, Asano Y, Muto A, Kawakami K, Suzuki Y, Kojima D, Fukada Y (2019) Six6 and Six7 coordinately regulate expression of middle-wavelength opsins in zebrafish. Proceedings of the National Academy of Sciences 116:4651–4660.

Palacios AG, Goldsmith TH, Bernard GD (1996) Sensitivity of cones from a cyprinid fish (Danio aequipinnatus) to ultraviolet and visible light. Visual Neuroscience 13:411–421.

Raine J, Hawryshyn C (2009) Changes in thyroid hormone reception precede SWS1 opsin downregulation in trout retina. Journal of Experimental Biology 212:2781–2788.

Raymond PA, Barthel LK, Rounsifer ME, Sullivan SA, Knight JK (1993) Expression of rod and cone visual pigments in goldfish and zebrafish: a rhodopsin-like gene is expressed in cones. Neuron 10:1161–1174.

Roberts Mr, Srinivas M, Forrest D, Morreale de Escobar G, Reh TA (2006) Making the gradient: thyroid hormone regulates cone opsin expression in the developing mouse retina. Proceedings of the National Academy of Sciences 103:6218–6223.

Saszik S, Bilotta J, Givin CM (1999) ERG assessment of zebrafish retinal development. Vis Neurosci 16:881–888.

Shen Y-c, Raymond PA (2004) Zebrafish cone-rod (crx) homeobox gene promotes retinogenesis. Developmental biology 269:237–251.

Sillman AJ, Ito H, Tomita T (1969) Studies on the mass receptor potential of the isolated frog retina: I. General properties of the response. Vision Research 9:1435–1442.

Suzuki SC, Bleckert A, Williams PR, Takechi M, Kawamura S, Wong RO (2013) Cone photoreceptor types in zebrafish are generated by symmetric terminal divisions of dedicated precursors. Proceedings of the National Academy of Sciences of the United States of America 110:15109–15114.

Swaroop A, Kim D, Forrest D (2010) Transcriptional regulation of photoreceptor development and homeostasis in the mammalian retina. Nature reviews Neuroscience 11:563–576.

Takechi M, Kawamura S (2005) Temporal and spatial changes in the expression pattern of multiple red and green subtype opsin genes during zebrafish development. Journal of Experimental Biology 208:1337–1345.

Takechi M, Hamaoka T, Kawamura S (2003) Fluorescence visualization of ultraviolet-sensitive cone photoreceptor development in living zebrafish. FEBS Letters 553:90–94.

Takechi M, Seno S, Kawamura S (2008) Identification of cis-Acting Elements Repressing Blue Opsin Expression in Zebrafish UV Cones and Pineal Cells. Journal of Biological Chemistry 283:31625–31632.

Terakita A (2005) The opsins. Genome biology 6:213.

Tsujimura T, Hosoya T, Kawamura S (2010) A single enhancer regulating the differential expression of duplicated red-sensitive opsin genes in zebrafish. PLoS genetics 6:e1001245.

Weiss AH, Kelly JP, Bisset D, Deeb SS (2012) Reduced L-and M-and increased S-cone functions in an infant with thyroid hormone resistance due to mutations in the THRβ2 gene. Ophthalmic genetics 33:187–195.

Wong KY, Gray J, Hayward CJ, Adolph AR, Dowling JE (2004) Glutamatergic mechanisms in the outer retina of larval zebrafish: analysis of electroretinogram b-and d-waves using a novel preparation. Zebrafish 1:121–131.

Yin J, Brocher J, Linder B, Hirmer A, Sundaramurthi H, Fischer U, Winkler C (2012) The 1D4 antibody labels outer segments of long double cone but not rod photoreceptors in zebrafish. Investigative ophthalmology & visual science 53:4943–4951.

Yoshimatsu T, Williams PR, D’Orazi FD, Suzuki SC, Fadool JM, Allison WT, Raymond PA, Wong RO (2014) Transmission from the dominant input shapes the stereotypic ratio of photoreceptor inputs onto horizontal cells. Nature communications 5.

